# Induction of NASH in the *Nwd1^−/−^* mouse liver via SERCA2-dependent endoplasmic reticulum stress

**DOI:** 10.1101/2024.01.26.577307

**Authors:** Seiya Yamada, Kazuhiko Nakadate, Tomoya Mizukoshi, Kiyoharu Kawakami, Ryosuke Kobayashi, Takuro Horii, Izuho Hatada, Shin-ichi Sakakibara

**Affiliations:** Laboratory for Molecular Neurobiology, Faculty of Human Sciences, Waseda University, 2-579-15 Mikajima, Tokorozawa, Saitama 359-1192, Japan; Department of Basic Biology, Educational and Research Center for Pharmacy, Meiji Pharmaceutical University, 2-522-1 Noshio, Kiyose-shi, Tokyo, 204-8588, Japan; Laboratory of Genome Science, Biosignal Genome Resource Center, Institute for Molecular and Cellular Regulation, Gunma University, 3-39-15 Showa-Machi, Maebashi, Gunma, 371-8512, Japan; Viral Vector Core, Gunma University Initiative for Advanced Research (GIAR), Gunma, 371-8511, Japan

**Keywords:** NASH, Nwd1, SERCA2, liver, lipid droplet, ER, ER stress, vacuole

## Abstract

The endoplasmic reticulum (ER) stores Ca^2+^ and plays crucial roles in protein folding, lipid transfer, and it’s perturbations trigger an ER stress. In the liver, chronic ER stress is involved in the pathogenesis of nonalcoholic fatty liver disease (NAFLD) and nonalcoholic steatohepatitis (NASH). Previous studies revealed that dysfunction of sarco/endoplasmic reticulum calcium ATPase (SERCA2), a key regulator of Ca^2+^ transport from the cytosol to the ER, is associated with the induction of ER stress and lipid droplet formation. We previously identified NACHT and WD repeat domain-containing protein 1 (Nwd1), which is localized in the ER and mitochondria. However, the physiological significance of Nwd1 outside the central nervous system remains unclear. In this study, we revealed that *Nwd1* knockout mice exhibited pathological manifestations comparable to NASH. Nwd1 interacts with SERCA2 near ER membranes. *Nwd1^−/−^* livers exhibited reduced SERCA2 ATPase activity and a smaller Ca^2+^ pool in the ER, leading to an exacerbated state of ER stress. These findings highlight the importance of SERCA2 activity mediated by Nwd1 in the pathogenesis of NASH.

**Highlights:**

- *Nwd1^−/−^* mice exhibited NASH-like liver steatosis.
- Elevated ER stress, fibrosis, and pyroptosis were observed in *Nwd1^−/−^* livers.
- Nwd1 interacts with SERCA2, an ER membrane Ca2^+ pump.^
- *Nwd1^−/−^* livers exhibited reduced SERCA2 activity and smaller Ca2+ pools in the ER.

## Introduction

Nonalcoholic fatty liver disease (NAFLD) and nonalcoholic steatohepatitis (NASH) are liver diseases that progress without symptoms, and they are linked to global public health problems. Characterized by the accumulation of lipid droplets in the liver, NASH evolves from steatosis to inflammation and cellular damage, including cell death, often culminating in fibrosis, cirrhosis, and eventually hepatocellular carcinoma [1]. The etiology of hepatic lipid accumulation involves various factors, including the excessive uptake or synthesis of fatty acids, defective transport of very low-density lipoprotein (VLDL), and impaired beta-oxidation of fatty acids [2,3]. Furthermore, spatially controlled hepatic zonation is crucial for physiological liver function, encompassing the metabolism of various endogenous products and xenobiotics [4–6]. A comprehensive understanding of the intricate interplay of cellular processes in NASH pathogenesis is imperative for developing targeted therapeutic interventions. Although several clinical trials of pharmacotherapies for NASH treatment are ongoing [1], no effective treatment has been developed.

NAFLD and NASH are considered polygenic disorders entailing a diverse array of genes, including unidentified genes. However, the individual contributions of these genes to the pathogenesis of these diseases are unclear. One crucial aspect of NASH pathology is the dysregulation of endoplasmic reticulum (ER) homeostasis [7,8]. The ER stores Ca^2+^ and plays crucial roles in protein folding, lipid transfer, and organelle dynamics regulation. The accumulation of unfolded or misfolded proteins in the ER activates a series of homeostatic responses collectively termed ER stress, and chronic ER stress contributes to the pathogenesis of NAFLD and NASH, including liver steatosis and lipid droplet formation [9,10]. The ER-localized transmembrane protein BSCL2/SEIPIN, acting at ER–lipid droplet contact sites, facilitates the incorporation of protein and lipid cargo into growing lipid droplets and helps connect newly formed lipid droplets to the ER and stabilize ER–lipid droplet contacts [11]. Recent studies suggested that the abnormal regulation of Ca^2+^ transport proteins is the basis of NAFLD/NASH [12,13]. The expression of type II inositol 1,4,5-trisphosphate receptor, the principal Ca^2+^-release channel in hepatocytes, was dramatically decreased in biopsied livers from patients with steatosis and NASH [14]. Consistently, dysfunction of sarco/endoplasmic reticulum calcium ATPase (SERCA2), a key regulator of Ca^2+^ transport from the cytosol to the ER, induces ER stress and lipid droplet formation [15]. Conversely, SERCA2 overexpression in obese mice reduced lipid synthesis gene expression and triglyceride content, emphasizing its significance in dysregulated lipid homeostasis and suggesting that SERCA2 could be targeted to treat NASH [16]. However, the molecular mechanism by which SERCA2 acquires its proper function in ER remains unclear.

Our prior research identified the NACHT and WD repeat domain-containing protein 1 (Nwd1) gene, a novel member of the signal transduction ATPases with numerous domains (STAND) protein superfamily [17,18]. Nwd1 is localized in the ER and mitochondria in neural stem/progenitor cells (NSPCs), and it is essential for normal brain development via interactions with the purine-synthesizing enzyme PAICS to regulate purinosome assembly in NSPCs [17,18]. Nwd1 is a multi-domain protein; i.e., it has a NACHT domain in the central region that is predicted to have nucleoside triphosphatase activity, an effector domain at the N-terminus that binds to PAICS, and a cluster of WD40 repeats at the C-terminus that is involved in interactions with other proteins or ligands. Based on its domain structure, Nwd1 is classified into the STAND superfamily, which is known to be involved in the formation of large protein complexes required for various signaling cascades such as apoptosis (apoptosome) and inflammation (inflammasome) and is conserved across species, including zebrafish, mice, rats, monkeys, and humans [17,19,20]. We previously demonstrated that Nwd1 is predominantly expressed in the liver in addition to the brain [17,19,20]; however, the physiological role of Nwd1 in nonneural tissues remains unknown.

In this study, we revealed that the livers of *Nwd1^−/−^*mice exhibit pathological manifestations resembling NASH, implying that *Nwd1* plays a role in modulating liver homeostasis by regulating the Ca^2+^ pump function of SERCA2. These findings could provide new clues to understanding the pathogenesis of ER stress-induced NASH.

## Materials and methods

### Animals

C57BL/6J mice purchased from Japan SLC, Inc. (Shizuoka, Japan) were housed under temperature- and humidity-controlled conditions on a 12-h/12-h light/dark cycle with *ad libitum* access to food and water. The day of birth was designated as P0.

### Generation of Nwd1*^−/−^* mice

All protocols were approved by the committees on the ethics of animal experiments at Gunma University and Waseda University. *Nwd1^flox^*mice were generated via electroporation, as previously reported with a partial modification [21]. Corresponding to the target DNA sequences (Nwd1L3, AATGGGACAGTCACTTGGGA; Nwd1R1, ACAGAACCTGGATTTTGTGG), donor single-stranded oligodeoxynucleotides (ssODNs) with 5′- and 3′-homology arms flanking loxP and a restriction site were designed (Nwd1L3loxpAS, cacacacacaTGCATGTGCTCACACATCCTAACCTGCCATGTGGGCTTTTGCAAGCCATCATAACTTCGTATAATGTATGCTATACGAAGTTATGGATCCCCAAGTGACTGTCCCATTTGTTAAACTTCACAGTCTGTGTCCCTCCGAACATGTTTCCCA; Nwd1R1loxP, GCCACATGCATGCTGCTTCCTTCCTTTTTTGTAGGGATTTGGACAGAACCTGGATTTTGTATAACTTCGTATAATGTATGCTATACGAAGTTATAAGCTTGGAGGTTAGTTATGTGTTGTGTGATTGAGCCTGGGCCCCAGACCtgcatgtagcctttgt; Fig. 1A). Pre-annealed crRNA (Alt-R CRISPR–Cas9 crRNA, Integrated DNA Technologies [IDT], Coralville, IA, USA])/tracrRNA (Alt-R CRISPR–Cas9 tracrRNA, IDT; 3 μM), recombinant Cas9 protein (100 ng/μl; GeneArt Platinum Cas9 Nuclease, Thermo Fisher Scientific, Waltham, MA, USA), and ssODNs (400 ng/μl; Ultramer, IDT) in Opti-MEM I (Thermo Fisher Scientific) were used for electroporation. First, a left loxP site was introduced into intron 4 of *Nwd1* by electroporation using B6D2F1-derived zygotes, and then the edited embryos were transferred to the oviduct of pseudopregnant ICR female to obtain *Nwd1* left loxPed males. Next, *Nwd1^flox^* mice were generated by introducing a right loxP site into intron 5 of *Nwd1* using left loxPed male-derived zygotes. Floxed alleles were confirmed by sequencing PCR products using the following primer set: 5′-CAGGTGTGATATGTAAATACTTCTTTG-3′ (F2) and 5′-CAGGCCTAACAGAGCCAGAC-3′ (R2). We further converted the floxed allele to a null allele (*Nwd1^−^*) by crossing *Nwd1^flox^* heterozygous mice with CAG-Cre driver mice (B6.Cg-Tg[CAG-Cre]CZ-MO2Osb, RBRC01828; RIKEN BRC, Tokyo, Japan) harboring the CAG-promoter-driven Cre recombinase transgene [22]. Exon 5, which encodes the NACHT domain, was eliminated by Cre/loxP-mediated excision. The genotyping of floxed and null alleles was performed by genomic PCR using the following primer sets: 5′-TCACACATCCTAACCTGCCA-3′ (F1) and 5′-AGTAGGCCAAGCTCGATCTC-3′ (R1) for the floxed allele and F1 and R2 primers for the null allele.

**Fig. 1.**
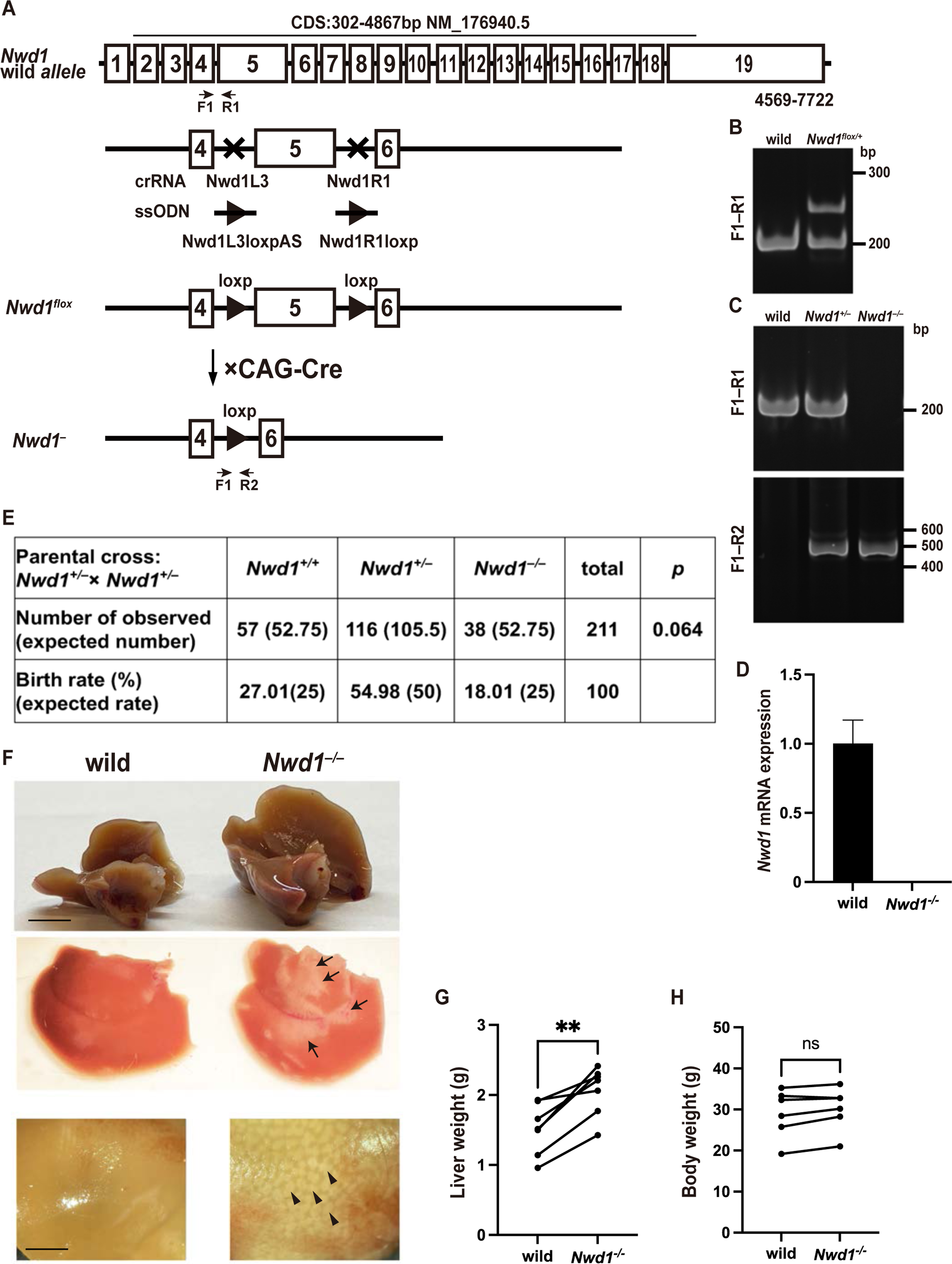
Generation of *Nwd1* knockout mice. (A) Schematic diagram of the *Nwd1* wild-type allele (top) and the conditional allele of *Nwd1*^flox^ (middle), in which LoxP sequences were inserted into the introns on both sides (crRNA targeting sites: Nwd1L3 and Nwd1R1) of exon 5 *via* two rounds of genomic editing using ssODNs (Nwd1L3loxpAS and Nwd1R1loxP). Nwd1^flox^ mice were crossed with CAG-Cre transgenic mice harboring CAG promoter-driven Cre recombinase to generate the *Nwd1*^−^ null allele (bottom). Upon elimination of the floxed *Nwd1* critical exon (exon 5), this allele generates a truncated RNA lacking the NACHT domain of the open-reading frame of *Nwd1*. (B) PCR genotyping of wild-type or *Nwd1*^flox/+^ tail genomic DNA. The specific primers (primer pair F1‒R1 depicted in A) were used to detect wild-type (189-bp amplicon) and *Nwd1*^flox^ (229-bp amplicon) alleles. PCR products were separated by 5% SDS-PAGE. (C) PCR genotyping of wild-type, *Nwd1*^+/−^, and *Nwd1*^−/−^ mice. Specific primers (primer pairs F1‒R1 and F2‒R2 depicted in A) were used to detect wild-type (189-bp amplicon) and *Nwd1*^−^ (431-bp amplicon) alleles. PCR products were separated by 5% SDS-PAGE. (D) Loss of *Nwd1* mRNA expression in *Nwd1*^−/−^ livers. The mRNA isolated from wild-type and *Nwd1*^−/−^ adult livers were subjected to qRT–PCR, and *Nwd1* expression was normalized to that of *β-actin* mRNA. (E) The table presents the genotypes and birth rates observed in 116 adult mice born from crosses between *Nwd1^+/−^*males and females. Numbers in parentheses represent the expected number of offspring and birth rate according to the Mendelian ratio. (F) Gross anatomical observation of the liver. *Nwd1*^−/−^ livers exhibited hepatomegaly (upper), fiber-like structures on the surface (middle, arrows), and numerous abnormal hepatic lobules that turned whitish (lower, arrowheads). (G, H) Liver (G) and body weight (H) of adult *Nwd1*^−/−^ mice and their wild-type littermates were compared by a paired *t*-test. ns, not significant; **, P < 0.01. Scale bars, 5 (F, upper) or 2 mm (F, lower).

### Cell culture and transfection

HEK293T and HeLa cells were cultured in Dulbecco’s Modified Eagle Medium (DMEM, FUJIFILM Wako Pure Chemical Corporation, Osaka, Japan) containing 10% fetal bovine serum (Biowest, Nuaillé, France), 1% penicillin/streptomycin (FUJIFILM Wako Pure Chemical), and 1% L-glutamine (FUJIFILM Wako Pure Chemical). Cells were seeded into poly-D-lysine–coated dishes (Merck Millipore, Burlington, MA, USA) and transfected with plasmid DNA and PEI MAX (Polysciences, Warrington, PA, USA) complexes (DNA to PEI MAX ratio, 1:3, w/w) formed in Opti-MEM (Thermo Fisher Scientific) via incubation for 15 min at 25°C. DNA complexes were added to cell cultures with Opti-MEM for 3 h, followed by cultivation with serum-containing DMEM.

### Electron microscopy

Anesthetized adult mice were perfused transcardially with saline solution, followed by a fixative containing 2% paraformaldehyde (PFA) and 2% glutaraldehyde (GA) in 0.1 M phosphate buffer (PB), pH 7.4. Liver tissue was dissected, cut into small pieces, washed several times with PB, and postfixed with 1% OsO_4_ (TAAB Laboratories, Aldermaston, UK) for 2 h. After dehydration with ethanol and replacement with propylene oxide, the sections were embedded in Epon-812 resin (TAAB Laboratories) and thermally polymerized. After trimming, the samples were cut into 70-nm-thick sections, electron-stained with uranium and lead, and observed and photographed using a transmission electron microscope (HT7800, Hitachi, Tokyo, Japan). For immunoelectron microscopy, HeLa cells expressing Flag-Nwd1 grown on glass coverslips were fixed with 4% PFA and 0.05% GA in 0.1 M PB (pH 7.4). The cells were washed with PB and treated with 30% sucrose in 0.1 M PB, and rapid freeze fractionation was performed using liquid nitrogen. Cells were blocked with normal goat serum for 2 h and incubated with anti-DDDDK-tag antibody for 2 h. After incubation with 1.4-nm gold-conjugated secondary antibody (Nanoprobes, Yaphank, NY, USA) for 1 h, the signals were gold-sensitized using a gold enhancing kit (Nanoprobes). After washing with H_2_O, cells were refixed with 1% OsO_4_ for 1 h, treated with ethanol and propylene oxide, and embedded in Epon-812 resin. Sections (70 nm thick) were electron-stained with uranium and lead and photographed using a transmission electron microscope (HT7800).

### Plasmids

pCAG-EGFP-Nwd1 was cloned as described previously (Yamada et al., 2020). pN3-3×Flag-Control was purchased from Addgene (Watertown, MA, USA, plasmid #107717, deposited by Dr. Suske) [23]. To construct 3×Flag-Nwd1 (hereafter called Flag-Nwd1), the full-length *Nwd1* cDNA was isolated using RT-PCR of total RNA from E12 embryonic and adult mice brains and then subcloned in-frame into the pN3-3×Flag-Control plasmid. *Nwd1* cDNA corresponding to the N-terminal or C-terminal region of the protein (accession number BC082552; 4–1026 bp for the N-terminus and 2578–4563 bp for the C-terminus) was subcloned into C-Flag-pcDNA3 (Addgene, plasmid # 20011, deposited by Dr. Smale) [24] and pN3-3×Flag-Control to construct Flag-Nwd1-N and 3×Flag-Nwd1-C (hereafter called Flag-Nwd1-C). To construct Halo-Nwd1, full-length Nwd1 cDNA was subcloned into the pFN21A-HaloTag-CMV Flexi-vector (Promega, Madison, WI, USA). To construct 3×Flag-SERCA2 (hereafter called Flag-SERCA2), the coding region of SERCA2b (Addgene, plasmid #75188, deposited by Dr. Lytton) [25] was subcloned in-frame into a pN3-3×Flag-Control vector. The deletion mutants 3×Flag-SERCA2-N (amino acids 1–787, hereafter called Flag-Flag-SERCA2-N) and 3×Flag-SERCA2-C (amino acids 788–1044, hereafter called Flag-SERCA2-C) were constructed from 3×Flag-SERCA2 using a Q5 Site-Directed Mutagenesis Kit (New England BioLabs, Ipswich, MA, USA). mCherry-Sec61B (Addgene, plasmid # 121160, deposited by Dr. Mayr) [26] was used for ER imaging.

### Primary antibodies

The following primary antibodies were used: anti-cleaved caspase-3 (Asp175) (rabbit polyclonal, #9661, RRID: AB_2341188, Cell Signaling Technology, Danvers, MA, USA; 1:400 for immunohistochemistry [IHC]), anti-cleaved caspase-1 (Ala317) p10 (rabbit polyclonal, #AF4022, RRID: AB_2845464, Affinity Biosciences, Cincinnati, OH, USA; 1:200 for IHC), anti-BSCL2/SEIPIN (rabbit polyclonal, #ab106793, RRID: AB_10974250, Abcam, Cambridge, UK; 1:200 for IHC, 1:1000 for western blotting [WB]), anti-KDEL (mouse monoclonal, #M181-3, RRID: AB_10693914, MBL, Woburn, MA, USA; 1:400 for IHC, 1:400 for immunocytochemistry [ICC], 1:5000 for WB), anti-LAMP1 (1D4B) (rat monoclonal, #sc-19992, RRID: AB_2134495, Santa Cruz Biotechnology, Dallas, TX, USA; 1:250 for IHC, 1:1000 for WB), anti-LC3A/B (D3U4C) XP (rabbit monoclonal, #12741, RRID: AB_2617131, Cell Signaling Technology; 1:200 for IHC, 1:2000 for WB), anti-Nwd1 (rabbit polyclonal generated via immunization with the recombinant mouse Nwd1 protein [18]; 1:400 for ICC), anti-SERCA2 ATPase Clone IID8 (mouse monoclonal, #NB300-529, RRID: AB_531361, Novus Biologicals, Centennial, CO, USA; 1:200 for ICC, 1:2500 for WB), anti-DDDDK-tag (mouse monoclonal, #M185-3, RRID: AB_11126775, MBL; 1:10000 for WB, 1:10000 for ICC), anti-ADP/ATP translocase 1/2 (ANT1/2, rabbit polyclonal, #17796-1-AP, RRID: AB_2190358, Proteintech, Rosemont, IL, USA; 1:2000 for WB), anti-voltage-dependent anion-selective channel protein 1 (VDAC1)/porin (mouse monoclonal, #ab14734, RRID: AB_443084, Abcam; 1:2000 for WB), anti-α-tubulin (rabbit polyclonal, #11224-1-AP, RRID: AB_2210206, Proteintech; 1:5000 for WB), anti-GRP78/BIP (rabbit polyclonal, #11587-1-AP, RRID: AB_2119855, Proteintech; 1:5000 for WB), anti-GRP94 (rabbit polyclonal, #14700-1-AP, RRID: AB_2233347, Proteintech; 1:5000 for WB), anti-activating transcription factor 6 (ATF6, rabbit polyclonal, #24169-1-AP, RRID: AB_2876891, Proteintech; 1:4000 for WB, 1:200 for IHC), anti-CD11B/integrin alpha M (rabbit polyclonal, #21851-1-AP, RRID: AB_2878927, Proteintech; 1:200 for IHC 1:2000 for WB), anti-RAB5A (rabbit polyclonal, #11947-1-AP, RRID: AB_2269388, Proteintech; 1:2000 for WB, 1:250 for IHC), and anti-RAB7A (rabbit polyclonal, #55469-1-AP, RRID: AB_11182831, Proteintech; 1:2000 for WB, 1:250 for IHC).

### Hematological and biochemical analysis of blood and plasma samples

Peripheral whole blood was collected and briefly mixed in Insepak II (Sekisui, Tokyo, Japan), followed by incubation at room temperature for 15–30 min. Subsequently, the mixture was centrifuged at 3500 rpm for 15 min, and the supernatant was collected as the plasma. Peripheral blood was collected using MiniCollect K2E K2EDTA tubes (Greiner Bio-One, Kremsmünster, Austria). Hematological and biochemical analyses were conducted by the BioSafety Research Center Inc. (Shizuoka, Japan).

### Oil red O staining, Sirius red staining, periodic acid-Schiff (PAS) staining, and Congo red staining

General staining of liver tissue sections was performed using an Oil Red O Stain Kit (for fat staining) (ScyTek Laboratories, Logan, UT, USA), Picro-Sirius Red Stain Kit (for collagen staining) (ScyTek Laboratories), Periodic Acid-Schiff (PAS) Diastase Stain Kit (for polysaccharides staining) (ScyTek Laboratories), and Amyloid Stain Kit (Congo Red) (for amyloid staining) (ScyTek Laboratories) according to the manufacturer’s protocol. Images were acquired using a DP72 CCD camera (Olympus, Tokyo, Japan) mounted on an ECLIPSE E800 (Nikon, Tokyo, Japan), BX53 (Olympus), or BZ-X800 microscope (Keyence, Osaka, Japan). Quantification of tissue images was performed using the hybrid cell count algorithm (Keyence).

### Immunostaining and TUNEL staining

Fluorescent immunostaining was performed as previously described [17]. Anesthetized adult mice were perfused transcardially with saline solution, followed by 4% PFA in 0.1 M PB (pH 7.4). Livers were dissected and postfixed in the same fixative overnight at 4°C. Fixed tissues were cryoprotect in 30% sucrose in 0.1 M PB (pH 7.4) overnight at 4°C and embedded in optimal cutting temperature compound (Sakura Finetek, Torrance, CA, USA). Frozen sections (14 μm thick) were collected on MAS-coated glass slides (Matsunami Glass, Bellingham, WA, USA). Sections were blocked for 2 h with 5% normal goat or donkey serum in 0.1% Triton X-100 in PBS (PBST), followed by incubation with primary antibodies in blocking buffer at 4°C overnight. After four washes with PBST, sections were incubated for 2 h with Alexa Fluor 488-, Alexa Fluor 555-, or Alexa Fluor 647-conjugated secondary antibodies (Thermo Fisher Scientific). After counterstaining with 0.7 nM Hoechst 33342 (Thermo Fisher Scientific) for the nucleus, 9 µM Bodipy (Thermo Fisher Scientific) for lipid droplets, or 75 µM Filipin III (Santa Cruz Biotechnology) for cytosol, sections were mounted and imaged using a confocal microscope (FV3000, Olympus). Filipin dye was used to delineate the entire cytoplasm, excluding vacuolar structures. To immunostain cultured cells, fixed cells were blocked for 1 h with 5% normal goat or donkey serum in PBST, followed by incubation with primary antibodies in blocking buffer for 1 h. After three washes with PBST, cells were incubated for 1 h with Alexa Fluor 488-, Alexa Fluor 555-, or Alexa Fluor 647-conjugated secondary antibodies. After counterstaining with 0.7 nM Hoechst 33342, the cells were mounted. To detect apoptosis, TUNEL staining was performed using the In Situ Cell Death Detection Kit, TMR-red (Roche Diagnostics, Basel, Switzerland) according to the manufacturer’s instructions.

### ER fraction and WB

The ER faction was obtained as previously described with some modifications [27,28]. Dissected fresh livers were homogenized in fraction buffer containing 10 mM HEPES (pH 7.4), 220 mM mannitol, 90 mM sucrose, and protease inhibitor cocktails (Complete mini, Merck Millipore) using Potter–Elvehjem homogenizers and then further homogenized via passage through a 27-gauge needle. The crude postnuclear supernatant was further clarified via centrifugation twice at 7000 × *g* for 10 min each and then at 10,000 × *g* for 10 min at 4°C. The pellet and supernatant were collected as the mitochondrial and cytosolic fractions, respectively. The cytosolic fraction was further separated into an ER-enriched pellet and cytosolic supernatant by ultracentrifugation (Optima LE-80K, Beckman Coulter, Brea, CA, USA) at 100,000 × *g* for 1 h at 4°C. The ER pellet was resuspended in the fraction buffer, and the total protein concentration was measured using the Bradford method (Bio-Rad, Hercules, CA, USA). For WB, the samples were treated with 2× sample buffer (125 mM Tris-HCl, pH 6.8, 4% SDS, 10% sucrose, and 0.01% bromophenol blue) for 1 h at 37℃, resolved by 8%–12% SDS-PAGE, and electroblotted onto an Immobilon-P membrane (Merck Millipore) using a semidry transfer apparatus. After blocking with 1%–5% skim milk in TBST (150 mM NaCl, 10 mM Tris-HCl, pH 7.4, 0.1% Tween 20), membranes were incubated with primary antibody for 1 h, followed by incubation with horseradish peroxidase (HRP)-conjugated secondary antibody (Cytiva [Marlborough, MA, USA] or Jackson ImmunoResearch [West Grove, PA, USA]). The signal was detected using Immobilon Western chemiluminescent HRP substrate (Merck Millipore). Blot images were captured using the Fusion Solo S system (Vilber Lourmat, Marne La Vallée, France) and quantified using the Quantification Software Module (Vilber Lourmat).

### Proteomic analysis and protein binding assay

HEK293T cells expressing Halo-Nwd1 were washed with ice-cold PBS and lysed in lysis buffer containing 50 mM Tris-HCl (pH 7.4), 150 mM NaCl, 1 mM EDTA, 1% NP-40, and protease inhibitor cocktail for 15 min at 4°C. The lysate was diluted 4-fold with Halo tag protein purification buffer containing 1× PBS, 1 mM DTT, and 0.005% NP-40 (Nacalai Tesque, Kyoto, Japan). The lysate was centrifuged at 15,000 rpm for 10 min at 4°C, and the supernatant was incubated with HaloLink Resin (Promega) overnight at 4°C. After brief centrifugation, the pelleted resin was washed four times with Halo tag protein purification buffer. To determine the proteins interacting with Halo-Nwd1, the pelleted resin was treated with 2× sample buffer for 1 h at 37°C and resolved by SDS-PAGE. The gel was stained using a Silver Stain MS Kit (FUJIFILM Wako Pure Chemical) according to the manufacturer’s instructions. Gel slices containing the protein bands were excised for proteomic analysis on a nanoLC-MS/MS system (Japan Proteomics Inc., Tokyo, Japan). Mass data acquisitions and peptide identification were performed using Mascot software. To verify the purification of recombinant halo-Nwd1 protein, the resin crosslinked with Halo-Nwd1 was incubated with HaloTEV protease (Promega) overnight at 4°C. Eluted Nwd1 protein was resolved by SDS-PAGE, followed by Coomassie brilliant blue (CBB) staining. For the protein binding assay, HEK293T cells expressing Halo-Nwd1 along with Flag-SERCA2, Flag-SERCA2-N, or Flag-SERCA2-C were lysed and purified using HaloLink Resin as previously described. Halo-Nwd1-binding proteins were treated with 2× sample buffer for 1 h at 37°C and resolved by SDS-PAGE, followed by WB using anti-Flag antibody.

### Quantitative PCR

Quantitative PCR was performed as previously described [29]. Total RNA prepared from adult mouse livers was reverse-transcribed using a PrimeScript II first-strand cDNA Synthesis Kit (Takara, Shiga, Japan) with a random hexamer primer. According to the manufacturer’s protocol, PCR was performed using TB Green Ex Taq II Mix (Takara) and the Thermal Cycler Dice Real-Time System (Takara).

### ATPase activity measurement

The ATPase activity of SERCA2 was measured using a previously described protocol with some modifications [15]. The ATPase activity of the ER fraction was measured using a QuantiChrom ATPase/GTPase assay kit (BioAssay Systems, Hayward, CA, USA) following the manufacturer’s instructions and quantified as colorimetric absorbance using a Nivo S luminometer (PerkinElmer, Shelton, MA, USA). The SERCA2 protein content in the ER fraction was determined by WB, and ATPase activity was normalized to that of total SERCA2 protein.

### Ca^2+^ measurement

The Ca^2+^ concentrations in ER and mitochondrial fractions were measured using the fluorescent indicator Fluo3. Each fraction was treated in buffer containing 100 mM KCl, 10 mM MOPS (pH 7.2), 0.01% Pluronic F-127, 2.6 µM Fluo3 (Dojindo, Kumamoto, Japan) in OptiPlate-96 Black plates (PerkinElmer). The fluorescent intensity (excitation: 495 nm, emission: 530 nm) was measured using a Nivo S luminometer. The total protein content of each fraction was determined using the Bradford method.

### Gene Ontology (GO) and Kyoto Encyclopedia of Genes and Genomes (KEGG) analysis

We used “DAVID Bioinformatics Resources” [30,31] to perform GO enrichment analysis (including biological process, cellular component, and molecular function) and KEGG pathway enrichment analysis [32].

### Statistical analyses

Statistical analyses were performed using Microsoft Excel version 16.77.1 (Microsoft, Redmond, WA, USA) and Prism version 9.5.0 (GraphPad, La Jolla, CA, USA). All numerical data are expressed as the mean ± SD. One-way analysis of variance (ANOVA) followed by Welch’s *t*-test with Holm–Bonferroni correction was used in multiple-group comparisons. Welch’s *t*-test was used to analyze between-group differences for the following variables: Hematoxylin and eosin (HE)^+^ normal hepatocytes, oil red O^+^ area, Sirius red^+^ fibrotic area, PAS^+^ area, the number of dead cells in the liver, the result of blood tests, the expression of ER stress-related proteins after tunicamycin (TM) treatment as determined by WB, relative ATPase activity, and relative Ca^2+^ concentrations. The paired *t*-test was used to compare liver and body weights and protein expression between littermates. The chi-squared test was used to compare the number of genotypes in the offspring of *Nwd1^+/−^*mating pairs. Significance was indicated by P < 0.05. We calculated the area of swollen hepatocytes, oil red O^+^ area, Sirius red^+^ fibrotic area, and PAS^+^ area using hybrid cell count software (Keyence). The numbers of lipid droplets and vacuolar structures in the liver were manually measured by polygon selection using cellSens imaging software (Olympus). Lipid droplets that partially filled the vacuole and detached from the vacuolar membrane were classified as type C vacuoles. The area of the vacuoles was measured as the area of the vacuolar structures minus that occupied by the lipid droplets.

## Results

### Generation of *Nwd1^−/−^* mice

The mouse Nwd1 gene was mapped to chromosome 8, and it consisted of 19 exons (Fig. 1A). To reveal the *in vivo* function of *Nwd1*, we generated *Nwd1*^−/−^ mice. *Nwd1^flox^*mice were generated by CRISPR-Cas9 genome editing using electroporation in zygotes. In total, 320 embryos were transferred to the recipient uteri. Of the 71 offspring born, five floxed mice were obtained. Conditional knockout of the *Nwd1^flox^* allele generated by genome editing was confirmed by PCR using genomic DNA (Fig. 1A, 1B). To obtain the mice harboring the *Nwd1^−^* null allele, heterozygous *Nwd1^flox/+^* mice were crossed with the CAG-Cre deleter mouse line, which ubiquitously expresses Cre recombinase, thereby allowing deletion of the loxP-flanked genomic region (Fig. 1A). The *Nwd1^−^* allele lacked the genomic region encompassing exon 5, a pivotal exon encoding the NACHT domain (Fig. 1A). Subsequent interbreeding of the heterozygous mutant mice yielded homozygous mutant (*Nwd1^−/−^*) pups. The accuracy of targeted disruption of *Nwd1* was confirmed through genomic sequencing and PCR (Fig. 1C). Quantitative reverse transcription (qRT)-PCR analysis confirmed the absence of *Nwd1* mRNA expression in *Nwd1^−/−^* mice (Fig. 1D). Genotyping of 211 adult offspring produced by breeding heterozygous *Nwd1^+/−^* mice revealed that the number of *Nwd1^−/−^*mice was 7% lower than expected from the Mendelian ratio (Fig. 1E). Histological examination demonstrated that most *Nwd1^−/−^* adult mice exhibited normal development and no evident abnormalities in any organs excluding the liver and lungs. The livers of *Nwd1^−/−^* mice exhibited an approximately 1.4-fold increase in liver size versus that in wild-type mice (wild-type, 1.5 ± 0.4; *Nwd1^−/−^*, 2.1 ± 0.4) despite no significant differences in body weight (wild-type, 29.0 ± 5.9; *Nwd1^−/−^*, 30.2 ± 5.2; Fig. 1F–H). Gross observation revealed abnormal speckled fatty-like patterns (Fig. 1F, arrowheads) and white fibrotic-like structures (Fig. 1F, arrows) on the surface of *Nwd1^−/−^* hepatic lobes.

### The *Nwd1^−/−^* liver exhibits NAFLD/NASH-like pathology

HE staining of *Nwd1^−/−^* livers revealed a striking increase in the number of enlarged and lucent hepatocytes with a lack of eosin staining compared to the findings in wild-type littermates (wild-type, 12.0% ± 3.1%; *Nwd1^−/−^*, 30.6% ± 3.5%; Fig. 2A, B).

**Fig. 2.**
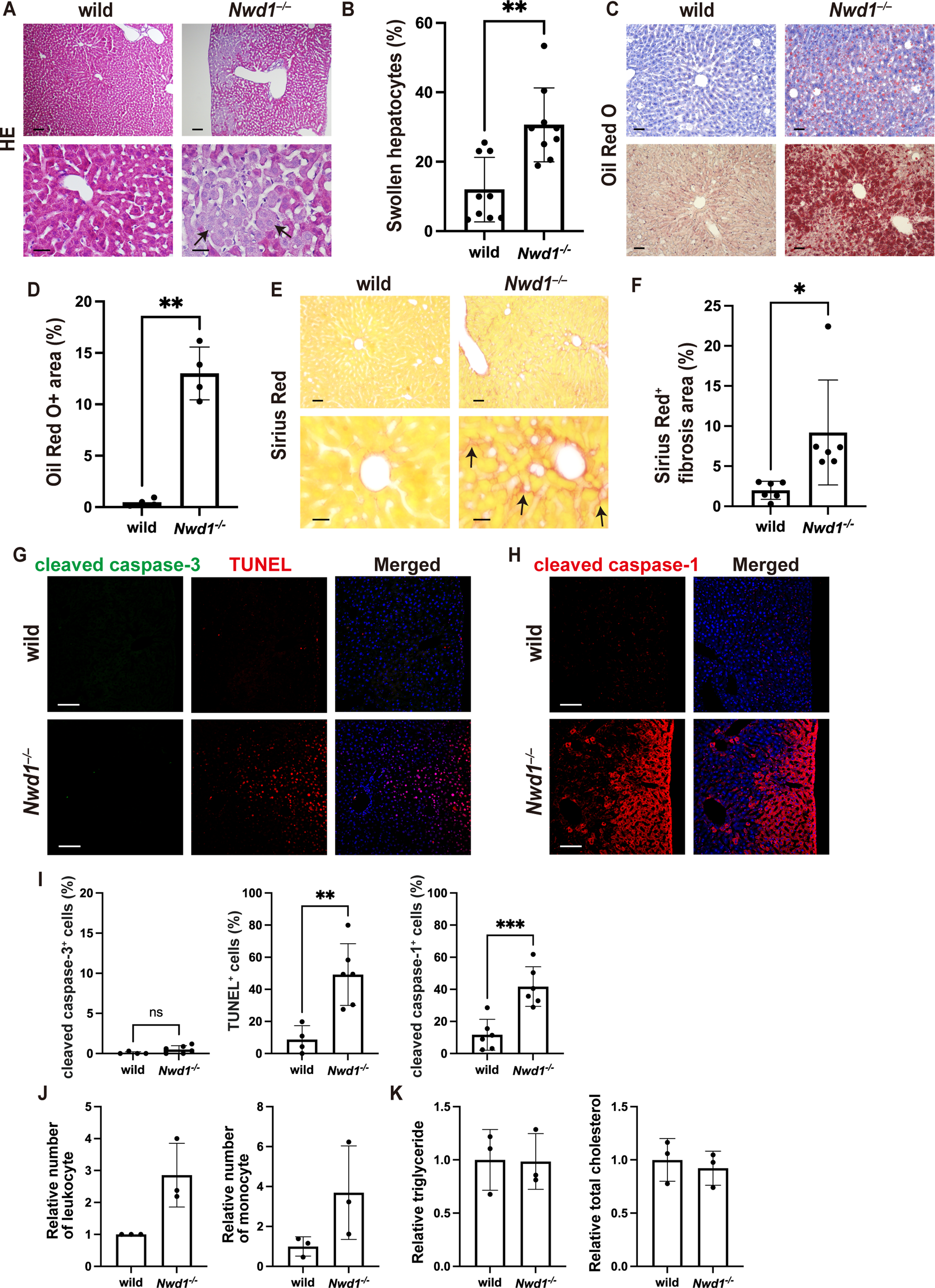
NASH-like phenotype in the adult *Nwd1*^−/−^ mouse liver. (A–I) Histological analyses of the adult mouse liver. HE (A, B), oil red O (C, D), and Sirius red (E, F) staining of *Nwd1*^−/−^ or wild-type livers. Nuclei were counterstained with hematoxylin (C, upper). Welch’s *t*-test was used to compare the number of swollen hepatocytes (%) (B), oil red O^+^ area (%) (D), and Sirius red^+^ fibrotic area (%) (F). *, P < 0.05; **, P < 0.01. Arrows denote swollen hepatocytes in (A) and Sirius red^+^ fibrotic pathology in (E). (G) Double staining with anti-cleaved caspase-3 antibody (*green*) and TUNEL (*red*). (H) Immunostaining using anti-cleaved caspase-1 (*red*) antibody. Nuclei were stained with Hoechst dye (*blue*). (I) Comparison of the number of cleaved caspase-3^+^ cells (%), TUNEL^+^ cells (%), and cleaved caspase-1^+^ cells (%) (Welch’s *t*-test). ns, not significant; **, P < 0.01; ***, P < 0.001. (J) Analysis of leukocyte and monocyte counts in peripheral blood (relative number). (K) Relative comparison of triglyceride and total cholesterol levels in serum. Scale bars, 10 (A [upper]), 40 (A [lower]) and (E [lower]), 50 (C and E [upper]), or 100 µm (G and H).

These swollen hepatocytes did not appear to have undergone programed cell death because their nuclei were not contracted or fragmented (Fig. 2A). We next performed oil red O staining on the liver tissues to detect the neutral lipid content in hepatocytes, finding that the size of the positively stained area was substantially increased in *Nwd1^−/−^*livers (wild-type, 0.5% ± 0.4%; *Nwd1^−/−^*, 13.0% ± 2.6%). Many lipid droplets, which are storage organelles for neutral lipids, accumulated in a zonate pattern around the central vein in the *Nwd1^−/−^*hepatic lobule (Fig. 2C, D). We further assessed the organization of collagen fibers in tissues using Sirius red staining. As presented in Fig. 2E and F, *Nwd1^−/−^* mice exhibited severe collagen fiber deposition in their livers (wild-type, 2.0% ± 1.1%; *Nwd1^−/−^*, 9.2% ± 6.5%), indicating that the loss of *Nwd1* resulted in abnormal fibrosis that resembled NASH. Congo red staining (Fig. S2A) illustrated that *Nwd1^−/−^* mouse livers did not accumulate amyloid fibrils, an extracellular deposition of amyloid proteins frequently observed in hepatic amyloidosis. PAS staining, which detects polysaccharides such as glycogen and mucosubstances such as glycoproteins and mucins, was significantly reduced in swollen *Nwd1^−/−^* hepatocytes (wild-type, 98.2% ± 0.15%; *Nwd1^−/−^*, 61.5% ± 18.6%; Figs. 2A and S1B, C). Because pretreatment with α-amylase abolished PAS staining in liver sections, insoluble glycogen was not accumulated in the *Nwd1^−/−^* mouse liver (Fig. S1B).

Based on the swelling of hepatocytes and detection of fibrosis, we explored the possibility of accelerated apoptosis in the *Nwd1^−/−^* liver using double staining with anti-cleaved caspase-3 antibody and TUNEL staining. As presented in Fig. 2G, although many TUNEL^+^ cells were observed in *Nwd1^−/−^* livers (wild-type, 8.7% ± 8.6%; *Nwd1^−/−^*, 49.2% ± 19.2%), few caspase-3^+^ apoptotic cells were found in these livers (wild-type, 0.1% ± 0.2%; *Nwd1^−/−^*, 0.5% ± 0.5%). TUNEL staining is known to detect different types of cell death [33], including pyroptosis, a form of programed necrotic cell death activated upon inflammation [34,35]. A much larger percentage of cells was positive for cleaved caspase-1, a pyroptosis marker, in *Nwd1^−/−^* mouse livers (wild-type, 11.7% ± 9.6%; *Nwd1^−/−^*, 41.7% ± 12.3%; Fig. 2H, I), indicating accelerated proptosis. The observation that *Nwd1^−/−^* hepatocytes lacked nuclear fragmentation, a morphological sign of apoptosis, was also consistent with the observation of inflammatory pyroptosis in *Nwd1^−/−^*livers. To evaluate the inflammatory response and lipid metabolism, hematological and biochemical analyses were performed using serum and peripheral whole blood. The total number of leukocytes was approximately 3-fold higher in *Nwd1^−/−^* mice than in wild-type mice (relative ratio = 2.85 ± 1.00; Fig. 2J). Although *Nwd1^−/−^* mice had increased counts of each type of leukocytes, including monocytes (relative ratio = 3.41 ± 4.29; Fig. 2J), lymphocytes, neutrophils, eosinophils, and basophils, the proportion of each cell type was unchanged (Fig. S2A, B). Lipid analysis revealed that the total content of triglycerides (relative ratio = 0.98 ± 0.26) and cholesterol (relative ratio = 0.92 ± 0.16) in the peripheral serum remained unchanged (Fig. 2K) despite the lipid accumulation in *Nwd1^−/−^* livers, suggesting impairment of the lipid droplet efflux in *Nwd1^−/−^* mice. The pathological phenotype of *Nwd1^−/−^* mice, including hepatomegaly, excessive lipid accumulation in hepatocytes, fibrosis, and inflammatory pyroptosis, closely resembled that of NASH.

In addition to the liver, we performed histological analysis of various organs. *Nwd1^−/−^*mouse lungs featured flattened alveoli (Fig. S3). We observed no abnormalities in the structure or tissue architecture of the heart, intestine, kidneys, skeletal muscle, spleen, and testes (Fig. S3).

### Accumulation of lipid droplets and ER-derived vacuoles in *Nwd1^−/−^*hepatocytes

Ultrastructure analysis using electron microscopy revealed that *Nwd1^−/−^*hepatocytes were occupied by numerous vesicular structures. These cytosolic vesicles varied in size, and they were divided into two types based on the inner structure: gray vesicles filled with electron-dense material and vacuoles that appeared vacant or contained small amounts of lipid (Fig. 3A, B). Ultrastructurally, both types of vesicles were surrounded by a single membrane (Fig. 3B). Based on these morphological characteristics, we concluded that the gray vesicles filled with electron-dense materials corresponded to the lipid droplets stained with oil red O. Notably, many tiny lipid droplets were evident in the wild-type liver sinusoid (Fig. 3A, arrows). However, we rarely observed lipid droplets in the *Nwd1^−/−^* sinusoid (Fig. 3B). To distinguish the vesicles, Bodipy staining of lipid droplets was performed in conjunction with cytoplasmic staining. Consistent with the result of electron microscopy, *Nwd1^−/−^* hepatocytes contained many vacant vacuoles in addition to lipid droplets filled with Bodipy^+^ triglycerides (Fig. 3C, D). We classified the vesicular structures based on Bodipy staining into three types (Fig. 3E): type A, a typical lipid droplet surrounded by a membrane; type B, a vacant vacuole lacking Bodipy^+^ lipids; and type C, a vacuole containing a small amount of Bodipy^+^ lipids, representing a transitional morphology between types A and B. The *Nwd1^−/−^* liver exhibited increases levels of all subtypes (Fig. 3E). Quantification of the areas occupied by lipid droplets and vacuoles in a single hepatocyte revealed a significant increase in both lipid droplets (wild-type, 1.98 ± 2.08 µm^2^; *Nwd1^−/−^*, 7.98 ± 6.96 µm^2^) and vacuolar structures (wild-type, 3.95 ± 2.46 µm^2^; *Nwd1^−/−^*, 12.18 ± 9.13 µm^2^) in *Nwd1^−/−^* mice (Fig. 3F, H). The numbers of giant lipid droplets larger than 2.5 μm^2^ (Fig. 3G) and vacuoles larger than 5 μm^2^ (Fig. 3I) were significantly increased in *Nwd1^−/−^* mice. Based on the increased size and number of lipid droplets, the absence of lipid droplets in the sinusoids (Fig. 3B), and the unchanged levels of total triglycerides and cholesterols in peripheral blood (Fig. 2K), we assumed that *Nwd1^−/−^* mice have defective lipid droplet efflux from the sinusoids, in addition to increased lipid synthesis in hepatocytes.

**Fig. 3.**
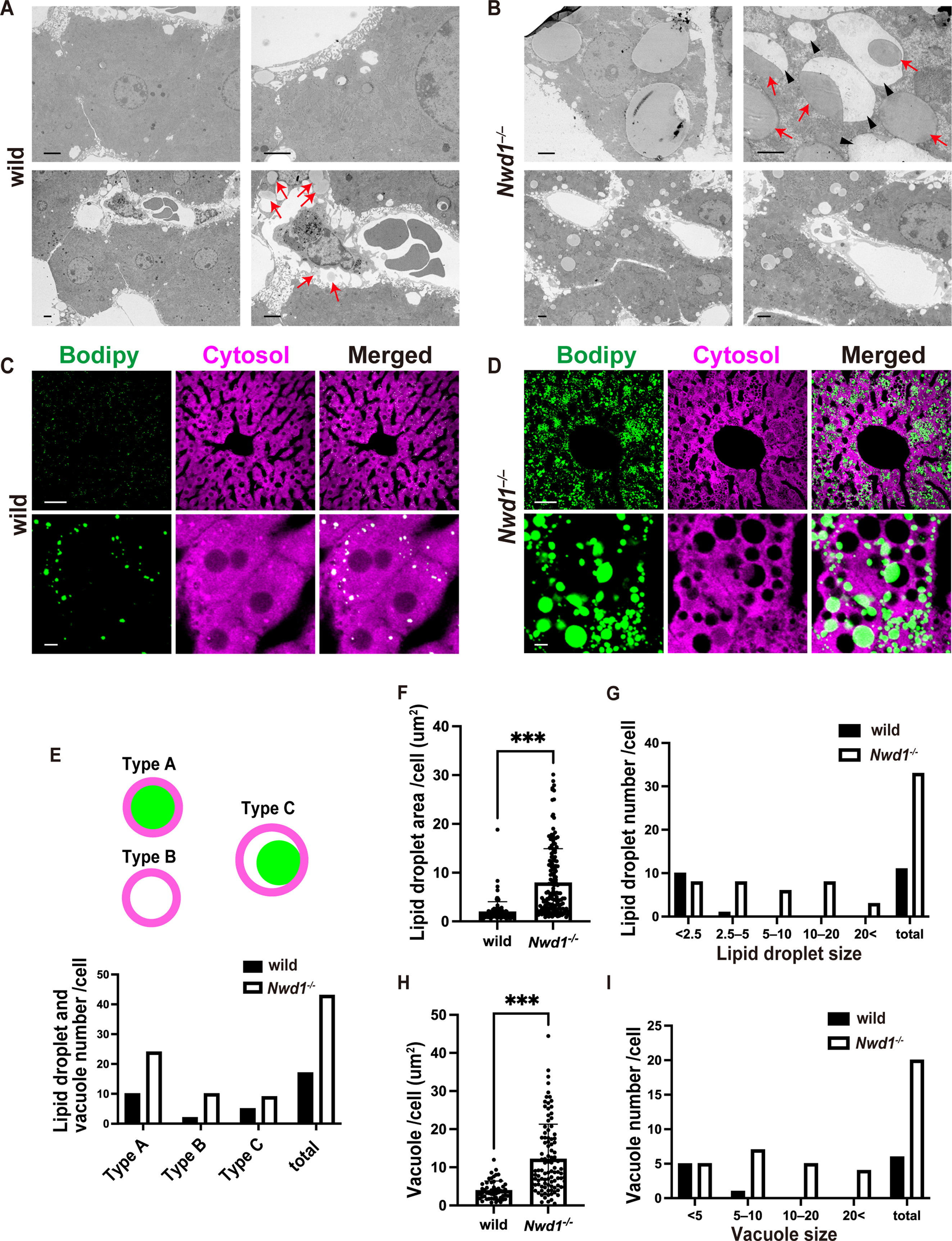
Accumulation of lipid droplets and vacuolar structures in *Nwd1*^−/−^ hepatocytes. (A, B) Electron microscopy of wild-type (A) and *Nwd1*^−/−^ (B) adult mouse liver. Arrows in (A) indicate the secretion of lipid droplets from hepatocytes into the wild-type liver sinusoid. (B) *Nwd1*^−/−^ hepatocytes contained numerous large vacuolar structures (black arrowheads) partially or entirely filled with electron-dense lipid droplets (red arrows). (C, D) Lipid droplet staining with Bodipy dye (*green*) in wild-type (C) and *Nwd1*^−/−^ livers (D). The cytosol (*magenta*) was counterstained. (E) Vesicular structures observed in *Nwd1*^−/−^ hepatocytes were classified into three types: type A, a typical lipid droplet surrounded by a membrane; type B, a vacant vacuole lacking Bodipy^+^ lipids; and type C, a lipid droplet partially or fully detached from the surrounding vacuolar membrane. The bars denote the distributions of types A, B, and C vesicles in wild-type and *Nwd1*^−/−^ hepatocytes. (E–I) Quantification of the area occupied by lipid droplets (F), the number and size of lipid droplets (G), the area occupied by vacuolar space (H), and the number of vacuoles (I) in wild-type and *Nwd1*^−/−^ hepatocytes. Welch’s *t*-test, ***, P < 0.001. Scale bars, 2 (A and B), 50 (C [upper] and D [upper]), or 5 µm (C [lower] and D [lower]).

From where do these empty vacuoles originate? Previous studies revealed that lipid droplet biogenesis is initiated in the ER, and ER-derived membrane structures enclose the lipid during the maturation of lipid droplets [11,15,36-38]. Cells lacking SEIPIN/BSCL2, a protein localized to ER–lipid droplet contact sites, had fewer ER– lipid droplet contact sites, and the transport of lipid droplets occurred independently of the ER [11]. Immunostaining for SEIPIN demonstrated that the *Nwd1^−/−^* liver exhibited normal SEIPIN^+^ zonation surrounding the central vein, and the area of SEIPIN^+^ zonation remained unchanged (Fig. 4A, B). Hepatocytes play distinct metabolic roles depending on their specific location along the lobular postcentral axis. This regional segregation of hepatocytes is called zonation. Consistently, the accumulation pattern of Bodipy^+^ lipid droplets within *Nwd1^−/−^* hepatic lobules appeared to be distributed as hepatic zonation surrounding a central vein (Fig. 4B). Immunostaining for the ER marker KDEL illustrated that many vacuoles in *Nwd1^−/−^* hepatocytes were surrounded by SEIPIN^+^ KDEL^+^ membranes, confirming that they were derived from the ER (Fig. 4C). Interestingly, RAB5, an early endosome marker, was frequently expressed in these ER-derived vacuoles (Fig. 4D). By contrast, RAB7, an early-to-middle endosome marker, exhibited little localization to the vacuoles (Fig. 4E). Moreover, the vacuolar structures did not express LAMP1 or LC3A/B, indicating that they are unrelated to the autophagy–lysosome–proteasome pathway (Figs. 4F, G and S4A, B).

**Fig. 4.**
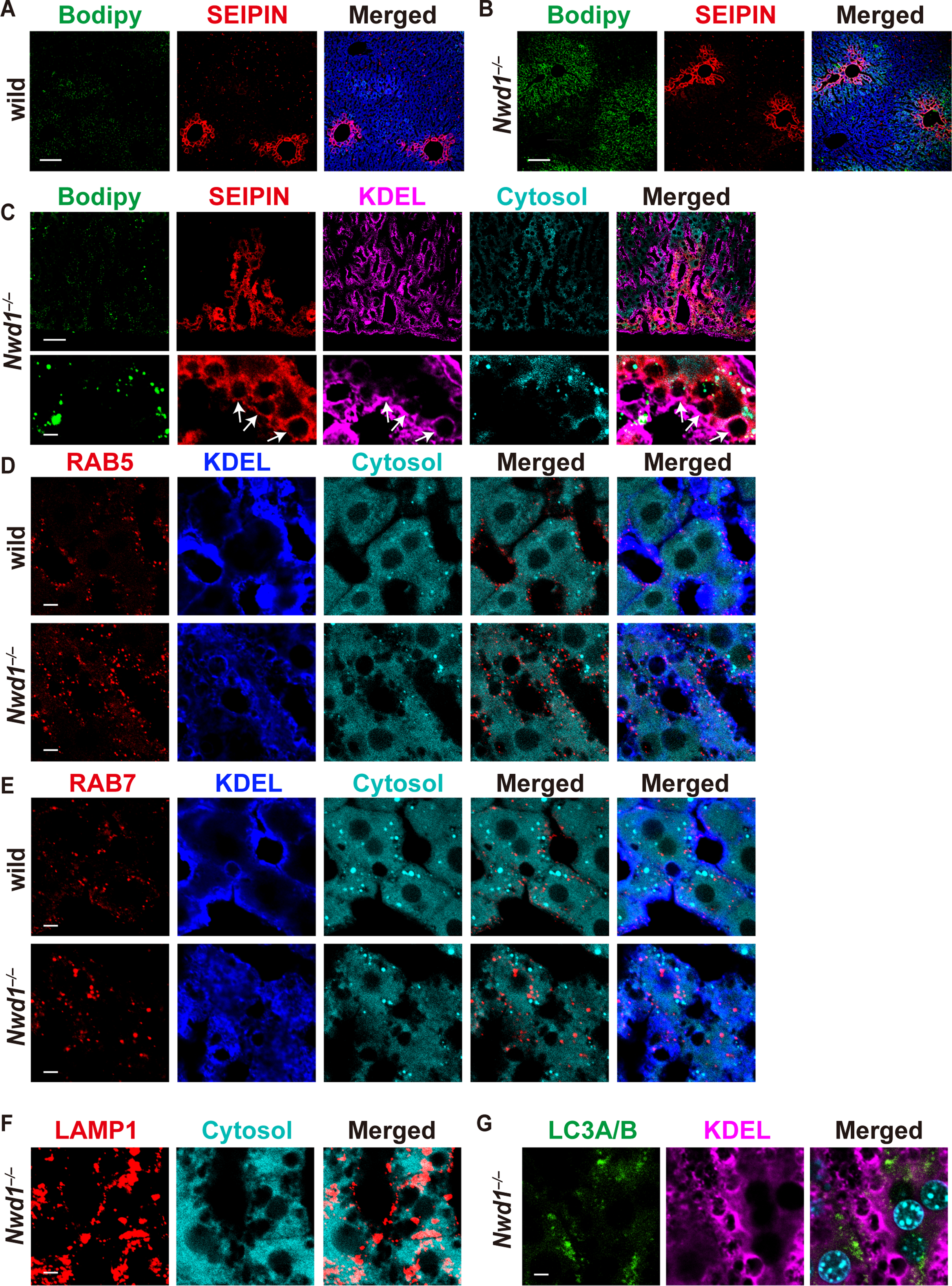
Lipid droplets and vacuolar structures of *Nwd1*^−/−^ hepatocytes were derived from the ER. (A, B) Lipid droplets (Bodipy, *green*) and SEIPIN (*red*) staining of wild-type (A) and *Nwd1*^−/−^ livers (B). Note the SEIPIN^+^ zonation surrounding the central vein in the *Nwd1*^−/−^ hepatic lobule. (C) Magnified view of the SEIPIN^+^ (*red*) area in *Nwd1*^−/−^ lobules simultaneously stained with Bodipy (*green*), anti-KDEL (*magenta*), and cytoplasm (*cyan*). Lower panels present a high-magnification view of the *Nwd1*^−/−^ hepatic cord. Many vacuolar structures were encapsulated by ER-derived membranes that were positive for SEIPIN and KDEL (arrows). (D, E) Magnified view of the hepatic cord immunostained with anti-RAB5 (D) or anti-RAB7 (E) together with anti-KDEL (*blue*) antibody and cytosol staining (*cyan*). (F, G) *Nwd1*^−/−^ hepatic cord immunostained with anti-LAMP1 (F) or anti-LC3A/B (*green*) and anti-KDEL (*magenta*) antibodies (G). The cytoplasm or nuclei were counterstained (*cyan*) (G). Scale bars, 100 (A and B), 50 (C [upper]), or 5 µm (C [lower], D, E, F, and G).

### Elevated ER stress in the *Nwd1^−/−^* liver

We examined the expression of proteins associated with ER and other intracellular organelles, including mitochondria, endosomes, and lysosomes. Notably, the ER membrane proteins SERCA2 and SEIPIN were upregulated in the *Nwd1^−/−^* liver compared to their wild-type expression (Fig. 5A, B). We also confirmed that the *Nwd1^−/−^* liver exhibited elevated expression of a 100-kDa KDEL protein (Fig. 5A, B). Conversely, the expression of ANT2 and VDAC, which are proteins of the inner and outer mitochondrial membranes, respectively, remained unchanged in the *Nwd1^−/−^* liver (Fig. 5A, B). RAB5 and LAMP1 expression was slightly increased in the *Nwd1^−/−^* liver, whereas RAB7 and LC3A/B expression remained unchanged (Fig. 5A, B). We concluded that many ER-related proteins were upregulated in the *Nwd1^−/−^*mouse liver.

**Fig. 5.**
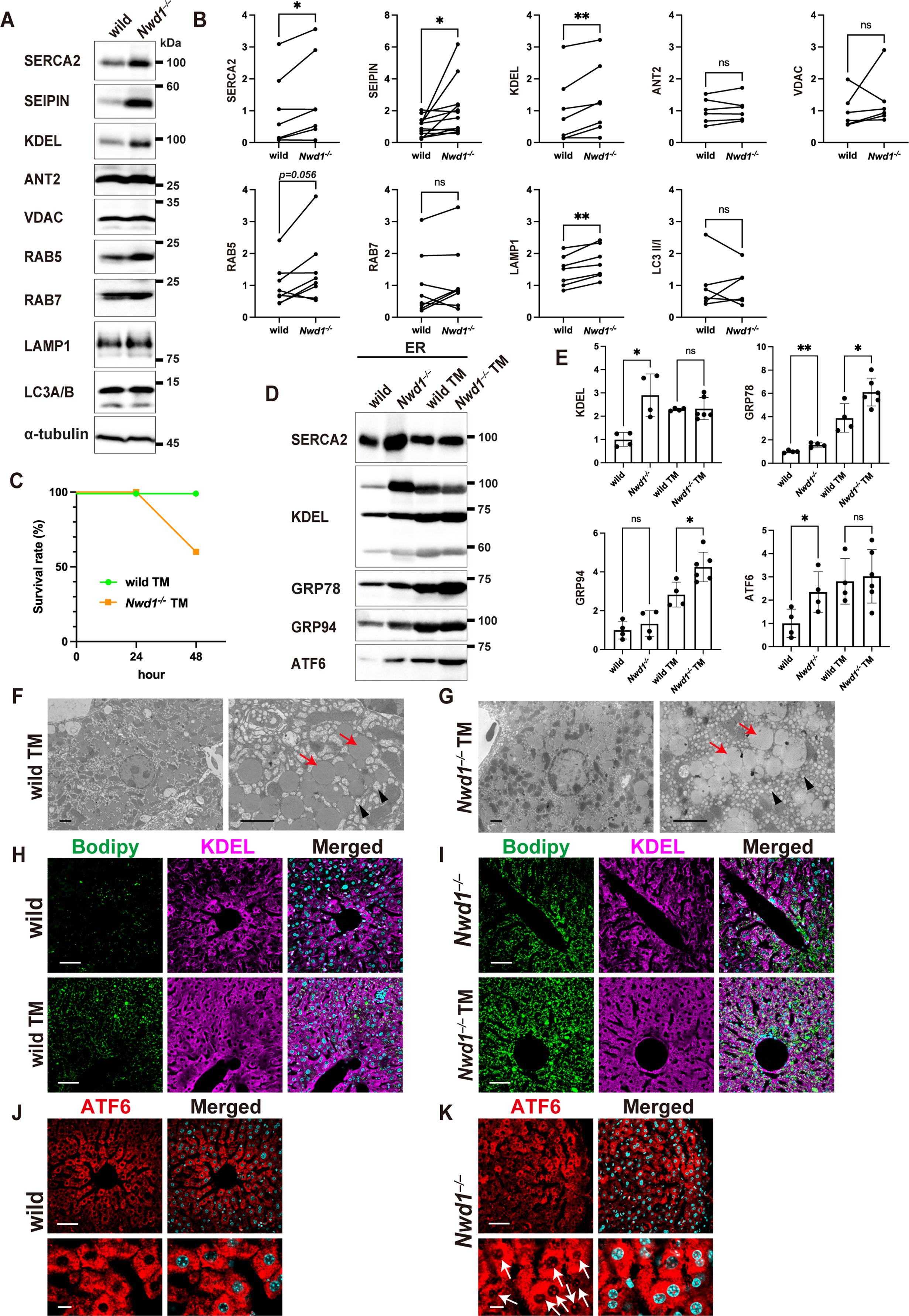
Increased ER stress in *Nwd1*^−/−^ livers. (A) WB of wild-type and *Nwd1*^−/−^ adult livers using the indicated antibodies. The blot was probed with anti-α-tubulin antibody to confirm equal loading. (B) Quantitative comparison of protein expression between an *Nwd1*^−/−^ mouse and its wild-type littermate (paired *t*-test). ns, not significant; *, P < 0.05; **, P < 0.01. (C) Survival curves of *Nwd1*^−/−^ and wild-type mice after TM treatment. (D) WB of ER stress-related proteins in ER fractions prepared from livers with or without TM treatment. (E) Quantitative comparison of the expression of each protein in (D). One-way ANOVA followed by Welch’s *t*-test with Holm–Bonferroni correction. ns, not significant; *, P < 0.05; **, P < 0.01. (F, G) Electron microscopy of *Nwd1*^−/−^ and wild-type hepatocytes treated with TM. A significantly increased number of small vacuoles (black arrowheads), in addition to large lipid droplets (red arrows), was observed in both wild-type and *Nwd1*^−/−^ livers after TM treatment. (H, I) Lipid (Bodipy, *green*) and ER (anti-KDEL, *magenta*) staining of wild-type and *Nwd1*^−/−^ livers with or without TM treatment. (J, K) ATF6 expression (*red*) in wild-type and *Nwd1*^−/−^ livers. Lower panels present magnified views. Arrows indicate nuclear translocation of ATF6, which occurred only in *Nwd1*^−/−^ livers. Nuclei were counterstained with Hoechst dye (*blue*). Scale bars, 2 (F and G), 50 (H, I, J [upper], and K [upper]), or 5 μm in (J [lower] and K [lower]).

Because the induction of ER stress is known to be accompanied by increased expression of ER-localized proteins such as KDEL proteins, it was speculated that ER stress was enhanced in the livers of *Nwd1^−/−^* mice. A previous study illustrated that tunicamycin (TM) treatment induced ER stress *in vivo* and the development of vacuolar structures in the mouse liver [39]. To assess the susceptibility of *Nwd1^−/−^*mice to ER stress, we intraperitoneally administered TM into wild-type and *Nwd1^−/−^*adult mice. Although wild-type mice survived up to 48 h after TM injection, 40% of *Nwd1^−/−^* mice died within 48 h (Fig. 5C), postulating that *Nwd1^−/−^*mice were more vulnerable to ER stress or that the basal level of ER stress was already elevated in these mice. To directly estimate whether ER stress is enhanced in the livers of *Nwd1^−/−^* mice, the ER fraction was prepared from wild-type and *Nwd1^−/−^* livers following treatment with or without TM, and the expression of ER stress-related proteins was quantified by WB (Fig. S4). ATF6, GRP78 (BiP), GRP94, and KDEL are known to be upregulated upon the induction of ER stress, and they have been used as markers of ER stress [40,41]. As presented in Figure 5D, TM administration to wild-type mice increased the expression of KDEL proteins harboring ER localization signal sequences, including GRP78 and GRP94, confirming that TM treatment induces ER stress in hepatocytes. At this time, the levels of these proteins were elevated in the ER fraction of *Nwd1^−/−^* livers regardless of TM treatment (KDEL: wild-type, 1.00 ± 0.30; *Nwd1^−/−^*, 2.90 ± 0.91; wild-type with TM treatment, 2.29 ± 0.06; *Nwd1^−/−^* with TM treatment, 233 ± 0.48; GRP78: wild-type, 1.00 ± 0.11; *Nwd1^−/−^*, 1.57 ± 0.19; wild-type with TM treatment, 3.88 ± 1.23; *Nwd1^−/−^* with TM treatment, 6.11 ± 1.12; and GRP94: wild-type, 1.00 ± 0.46; *Nwd1^−/−^*, 1.33 ± 0.67; wild-type with TM treatment, 2.83 ± 0.64; *Nwd1^−/−^* with TM treatment, 4.25 ± 0.76; Fig. 5D, E). Additionally, a significant increase in the expression of ATF6, an ER stress-responsive gene that encodes an ER-membrane-bound transcription factor activated by ER stress-induced proteolysis [39,41], was observed in the *Nwd1^−/−^* liver regardless of TM treatment (wild-type, 1.00 ± 0.61; *Nwd1^−/−^*, 2.35 ± 0.87; wild-type with TM treatment, 2.81 ± 0.98; *Nwd1^−/−^* with TM treatment, 3.02 ± 1.15; Fig. 5D, E). Electron microscopy revealed that TM-mediated ER stress drove the accumulation of large lipid droplets and vacuoles in the wild-type liver (Fig. 5F). This hepatic pathology closely resembled that in the *Nwd1^−/−^*liver without TM administration (Figs. 3B and 5G). Bodipy staining confirmed that TM-mediated ER stress accelerated the accumulation of lipid droplets in wild-type and *Nwd1^−/−^* livers (Fig. 5H, I). Immunostaining using ATF6 antibody illustrated that the nuclear translocation of ATF6 was induced in *Nwd1^−/−^*livers (Fig. 5K). ATF6 translocation was never detected in wild-type mice (Fig. 5J), indicating that ER stress is constitutively induced in the *Nwd1^−/−^* liver.

### Nwd1 interacts with ER-associated proteins, including SERCA2

To understand the molecular mechanism underlying ER stress induced by Nwd1 depletion, Nwd1-binding proteins were screened using the pulldown assay and proteomic analysis. Nwd1-interacting proteins were copurified with halo-tagged Nwd1 protein from HEK293T cell lysate (Fig. 6A). Proteomic analysis using nanoLC-MS/MS identified 107 proteins as candidate Nwd1-binding proteins, including SERCA2, the mitochondrial outer membrane proteins VDAC1 and sideroflexin-1, the endosome membrane protein RAB7A, and chaperone proteins including calnexin and heat shock 70 kDa protein 1A (Fig. 6B and Tables S1–S3). GO analysis for biological processes suggested a predominant role of Nwd1 in protein transport (Fig. 6C). According to the predicted distribution of the cellular components and the prediction of molecular function, many Nwd1-binding proteins were expected to localize to the ER membrane and various organelle membranes and function in protein binding (Fig. 6C). *In silico* pathway analysis using the KEGG database predicted that many Nwd1-binding proteins were involved in protein processing in the ER (Fig. 6C). Collectively, these analyses strongly predicted the involvement of Nwd1 in the membrane transport or processing of ER proteins.

**Fig. 6.**
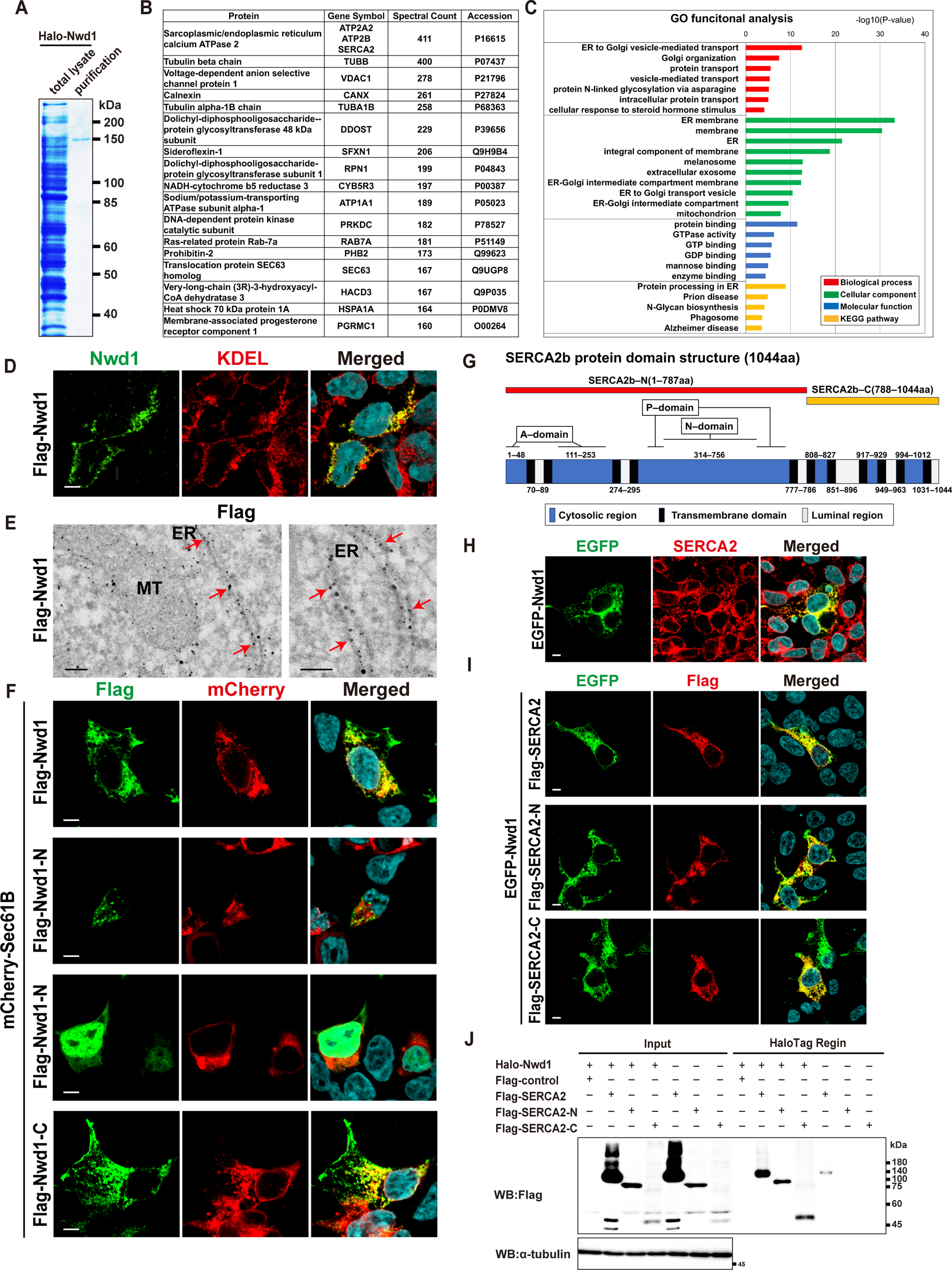
Nwd1 binds to SERCA2. (A) Purification of Halo-Nwd1 recombinant protein expressed in HEK293T cells. CBB staining of the SDS-PAGE gel of the cell lysate and purified protein. (B) Identification of Halo-Nwd1-binding proteins by nanoLC-MS/MS. The list of representative Nwd1-binding proteins presents the gene symbols, the spectral counts in proteomic analysis, and the UniProt protein database entry numbers. (C) GO enrichment and KEGG pathway analyses of Nwd1-binding proteins. (D) HEK293T cells expressing Flag-Nwd1. ICC with anti-Nwd1 (*green*) and anti-KDEL (*red*) antibodies revealing the co-localization of Nwd1 with the ER. (E) Immunoelectron microscopy revealing the localization of Flag-Nwd1 in the vicinity of the ER. HeLa cells were transfected with Flag-Nwd1 and immunostained with anti-Flag antibody (gold particle). Arrows indicate the Nwd1 signals on or near the ER membranes. MT, mitochondria. (F) Analysis of the Nwd1 region required for localization to the ER. HEK293T cells were transfected with mCherry-Sec61B and Flag-Nwd1, Flag-Nwd1-N, or Flag-Nwd1-C. Cells were stained with anti-Flag (*green*) antibody. (G) Structure and topology of mouse SERCA2b protein (1044 amino acids, accession number NP_733765.1). Blue boxes denote the cytosolic regions comprising the A (amino acids 1–43 and 124–235), N (amino acids 360–600), and P domains (amino acids 330–359 and 601–739), which are involved in ATPase activity. Black boxes denote 11 transmembrane domains, and white boxes denote the ER luminal regions. The red and yellow lines represent SERCA2b–N (amino acids 1– 787) and SERCA2b–C (amino acids 788–1044), the deletion mutants of SERCA2b produced in this study. (H) Co-localization of Nwd1 and SERCA2 in the ER. HEK293T cells were transfected with EGFP-Nwd1 and stained with anti-SERCA2 (*red*) antibody. (I) HEK293T transfected with EGFP-Nwd1 and Flag-SERCA2, Flag-SERCA2-N, or Flag-SERCA2-C were stained with anti-Flag (*green*) antibody. (J) *In vitro* binding assay of Nwd1 with SERCA2. Cell lysate of HEK293T cells expressing Halo-Nwd1 and Flag-SERCA2, Flag-SERCA2-N, or Flag-SERCA2-C were pulled down using HaloLink Resin, and the binding with Halo-Nwd1 was analyzed by WB using anti-Flag antibody. The blot probed with anti-α-tubulin antibody confirmed equal input of each lysate. Scale bars, 5 μm (D, F, H, and I) or 100 nm (E).

We examined whether Nwd1 is distributed in the ER. Double immunostaining demonstrated the co-localization of Nwd1 and KDEL in cultured HEK293T cells (Fig. 6D). To analyze the association between Nwd1 and ER at higher resolution, immunoelectron microscopy was performed using HeLa cells expressing Flag-Nwd1. As presented in Figure 6E, many Flag-Nwd1 signals were detected on and near the surface of the ER membrane but rarely in the lumen of the ER, confirming the close association of Nwd1 with the ER membrane. To identify the protein region responsible for ER localization, deletion mutants of Nwd1 were introduced into cultured cells together with mCherry-Sec61B to visualize the ER architecture [26,29]. The overlapping distribution of Flag-tagged full-length Nwd1 and mCherry-Sec61B verified the predominant localization of Nwd1 to the ER. Flag-Nwd1-N, which lacks the central NACHT domain and C-terminal WD40 repeat domain, localized to the cytoplasm or nucleus (Fig. 6F). Conversely, Flag-Nwd1-C, which only contains the WD40 repeat domain, was explicitly localized to the ER (Fig. 6F), indicating that the C-terminal WD40 repeat domain of Nwd1 is crucial for its ER localization and thus for interactions with ER proteins. Generally, the WD peptide motif folds into a β-propeller structure, and proteins containing WD40 repeats are believed to serve as platforms for assembling protein complexes or as mediators of transient interactions between other proteins [42].

Among the ER proteins identified as potential Nwd1-binding proteins, we focused on SERCA2b, which had the highest spectral count in the pulldown assay (Fig. 6B). SERCA2 is an ATPase pump embedded in the ER membrane that transports Ca^2+^ from the cytosol to the ER lumen at the expense of ATP hydrolysis [15]. Immunostaining revealed that the intracellular distribution of Nwd1 protein was identical to that of endogenous and exogenous SERCA2 protein (Fig. 6H). The protein topology of SERCA2b includes a cytoplasmic region comprising an actuator (A) domain (amino acids 1–43 and 124–235), a nucleotide-binding (N) domain (amino acids 360–600), a phosphorylation (P) domain (amino acids 330–359 and 601–739), and 11 transmembrane domains [43,44] (Fig. 6G). As previously described [45], we generated two deletion constructs of SERCA2b named SERCA2-N and SERCA2-C. SERCA2-N contains mainly the A, N, and P domains (amino acids 1–787), and SERCA2-C corresponds to the region containing the ER lumen and the short cytoplasmic region (amino acids 788–1044; Fig. 6G). SERCA2-N or SERCA2-C was introduced to HEK293T cells with Flag-Nwd1, and their intracellular distribution was monitored. As presented in Figure 6I, both SERCA2 deletion mutants localized to the ER similarly as full-length SERCA2. *In vitro* protein binding assays using Halo-Nwd1 and Flag-SERCA2 mutants demonstrated that Nwd1 binds to both the N-terminal region of SERCA2 outside the ER and the C-terminal region (Fig. 6J). These results indicate that Nwd1 recognizes a broad cytosolic region spanning the SERCA2 protein to regulate the trafficking, conformational folding, or embedding of SERCA2 into the ER membrane.

### Decreased ATPase activity of SERCA2 and smaller Ca^2+^ pools in the *Nwd1^−/−^* liver ER

How does the Nwd1–SERCA2 interaction contribute to ER stress or homeostasis? ER fractions were prepared from wild-type and *Nwd1^−/−^* livers, and SERCA2 ATPase activity, which regulates Ca^2+^ transport from the cytosol to the ER lumen, was measured. The results indicated that SERCA2 ATPase activity was significantly reduced by approximately 70% in the *Nwd1^−/−^* liver (wild-type, 1.00 ± 0.64; *Nwd1^−/−^*, 0.29 ± 0.09; Fig. 7A). Consistent with the decrease in ATPase activity, Ca^2+^ stores in the *Nwd1^−/−^* ER were depleted by approximately 40% (wild-type, 1.00 ± 0.09; *Nwd1^−/−^*, 0.63 ± 0.03; Fig. 7B). Contrarily, the amount of Ca^2+^ inside mitochondria did not differ between wild-type and *Nwd1^−/−^* livers (wild-type, 1.00 ± 0.30; *Nwd1^−/−^*, 0.92 ± 0.16; Fig. 7C). Considering the finding that SERCA2 protein expression was drastically increased by Nwd1 deficiency (Fig. 5), it was likely that the proportion of SERCA2 lacking pump activity is increased in *Nwd1^−/−^* livers. These results implied that Nwd1 is required for the folding or topology of SERCA2 on the ER membrane, enabling SERCA2 to acquire proper activity.

**Fig. 7.**
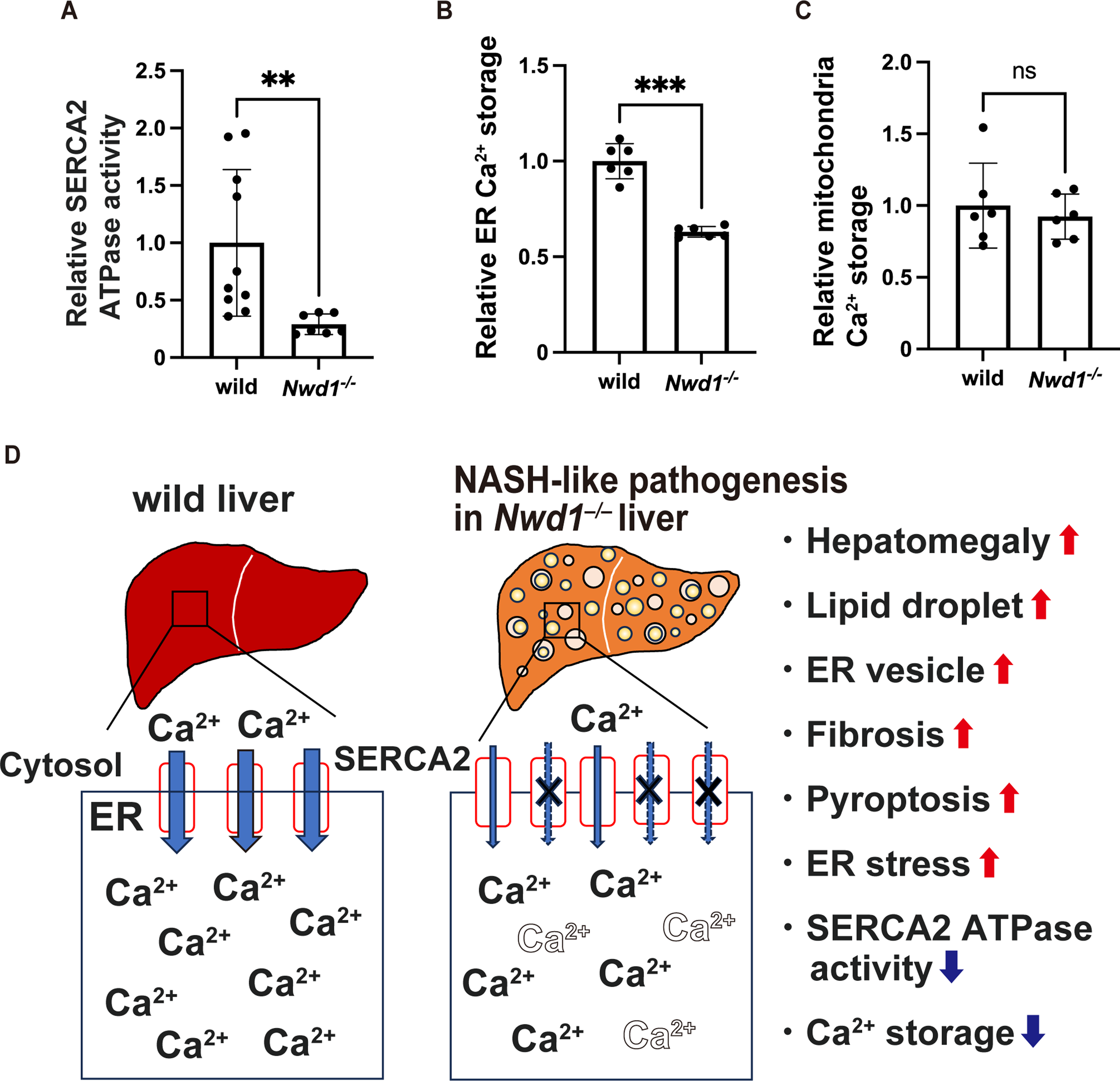
SERCA2 ATPase activity and ER Ca^2+^ storage are suppressed in the *Nwd1*^−/−^ liver. (A) SERCA2 ATPase activity in the ER fractions of wild-type and *Nwd1*^−/−^ livers. (B, C) Ca^2+^ storage in the ER (B) or mitochondrial (C) fractions of wild-type and *Nwd1*^−/−^ livers. Welch’s *t*-test, ns, not significant; **, P < 0.01; ***, P < 0.001. (D) Summary of NASH-like hepatic pathogenesis caused by *Nwd1* deficiency and a model for dysfunction of the SERCA2 Ca^2+^ pump in the *Nwd1*^−/−^ liver ER. Decreased activity of SERCA2 in the *Nwd1*^−/−^ liver induces ER stress by reducing Ca^2+^ stores in the ER, leading to diverse hepatic pathologies, including the accumulation of ER-derived vacuolar structures with lipid droplets, fibrosis, and pyroptosis. The ER membranes of *Nwd1*^−/−^ livers featured an increased population of SERCA2 with no or reduced pump activity (indicated by crosses and thin arrows), lowering Ca^2+^ stores inside the ER.

## Discussion

Previous studies illustrated that ER stress contributes to hepatic steatosis by inducing the accumulation of lipid droplets [8,9]. However, the mechanism by which ER stress in the liver influences the development of hepatic steatosis is unclear. We demonstrated that Nwd1 deficiency led to decreased Ca^2+^ storage in the ER *via* the reduction of SERCA2 ATPase activity, leading to ER stress accompanied by diverse hepatic pathologies, including the accumulation of lipid droplets and ER-derived vacuoles in hepatocytes, hepatomegaly, hepatic fibrosis, and caspase-1–dependent inflammatory pyroptosis along with an elevated number of leukocytes in peripheral blood (Fig. 7D). Many of the pathologies in *Nwd1*-deficient mice are comparable to those in NASH in humans. Quality control of newly synthesized proteins largely depends on the Ca^2+^ concentration in the ER lumen. For instance, the calcium-binding protein GRP78 plays a pivotal role in detecting the accumulation of misfolded proteins in the ER in cooperation with other ER transmembrane proteins such as ATF6, IRE1, and PERK [46,47]. Protein disulfide isomerases, a major group of ER-resident proteins, directly interact with Ca^2+^ to serve as molecular chaperones for protein synthesis and maturation. Therefore, dysregulation of Ca^2+^ homeostasis in the ER leads to the collapse of protein quality control, resulting in the induction of ER stress [46]. It is well established that ER stress associated with unfolded proteins causes the excessive accumulation of lipid droplets surrounded by a monolayer membrane of ER-derived phospholipids in the liver [7,9,41,48-50].

Despite the abundant accumulation of lipid droplets, no significant increases in serum triglyceride and total cholesterol levels were observed in the *Nwd1^−/−^* liver. Considering that lipid droplets in the sinusoid were rarely observed in the *Nwd1^−/−^* liver using electron microscopy, this NASH-like phenotype might be related to impaired hepatic fat clearance in addition to augmented *de novo* fat synthesis in the *Nwd1^−/−^*liver. Consistently, a previous report in mice treated with TM disclosed the induction of vacuolar structures in the liver and concomitant decreases in total cholesterol and triglyceride levels in plasma, presumably attributable to the suppression of VLDL emission from the liver [39]. Furthermore, recent studies highlighted the involvement of pyroptosis, a caspase-1–mediated programed cell death process, in the pathology of hepatic steatosis and NASH [51]. Pyroptosis is initiated by the activation of caspase-1 through inflammasome formation, leading to pore formation in the plasma membrane and the release of pro-inflammatory factors into the pericellular space. Specifically, the release of caspase-1–dependent inflammatory complexes in hepatocytes promotes liver fibrosis and the excessive accumulation of extracellular matrix proteins, including collagen [34]. Inhibiting pyroptosis has been reported to effectively reduce fat deposition in the liver and reduce liver inflammation in NASH [52]. Along the same lines, increased pyroptosis might have driven fibrosis in the *Nwd1^−/−^*liver. Deciphering the molecular functions of Nwd1 could shed light on the novel aspects of the etiology of diseases associated with increased ER stress and impaired Ca^2+^ homeostasis, such as NAFLD and NASH.

Nwd1 might play an essential role as a co-chaperone in modulating the activity or conformational regulation of SERCA2 on the ER membrane by binding to a specific domain of SERCA2. Immunoelectron microscopy revealed that Nwd1 localizes on the membrane or in close proximity to the cytoplasmic side of the ER, but it was rarely found within the ER. *In vitro* binding assays illustrated that Nwd1 recognizes and binds to a broad region of SERCA2, including cytosolic ATPase domains and the C-terminal short region, which is involved in the Ca^2+^ affinity of SERCA2b [53]. A previous study found that the topological fluctuation of SERCA2b is important for the transition of Ca^2+^ affinity [44]. Nwd1 might affect the Ca^2+^ affinity of SERCA2 by regulating its topology through binding to the C-terminal region, in addition to modulating ATPase activity through binding to the A, N, and P domains of SERCA2. The elevated expression of SERCA2 protein observed in *Nwd1^−/−^*livers might be attributable to a compensatory mechanism in response to the increased proportion of SERCA2 lacking proper pump activity.

Our previous study illustrated that Nwd1 regulates the formation of purinosomes during cerebral cortex development by interacting with PAICS, an enzyme required for *de novo* purine biosynthesis [18]. The purinosome is a macromolecular metabolic complex consisting of six enzymes that catalyze *de novo* purine biosynthesis and chaperone proteins necessary for the association of component proteins [54]. Nwd1 is also reported to interact with several chaperones in prostate cancer cell lines [55]. These facts led us to speculate that Nwd1 might function as a co-chaperone for protein folding, allowing target proteins, including SERCA2, to acquire appropriate activity. Elucidating the molecular machinery by which *Nwd1* is involved in hepatocellular pathology could provide clues to understanding the mechanisms of NASH pathogenesis and developing new therapeutic strategies for NASH associated with ER stress.

## Abbreviations

ER: endoplasmic reticulum
NAFLD: nonalcoholic fatty liver disease
NASH: nonalcoholic steatohepatitis
SERCA2: sarco/endoplasmic reticulum calcium ATPase
Nwd1: NACHT and WD repeat domain-containing protein 1
VLDL: very low-density lipoprotein
STAND: signal transduction ATPases with numerous domains
NSPCs: neural stem/progenitor cells
ssODNs: single-stranded oligodeoxynucleotides
DMEM: Dulbecco’s Modified Eagle Medium
PFA: paraformaldehyde
GA: glutaraldehyde
PB: phosphate buffer
PAS: periodic acid-Schiff
HRP: horseradish peroxidase
GO: Gene Ontology
KEGG: Kyoto Encyclopedia of Genes and Genomes
TM: tunicamycin
HE: hematoxylin and eosin
ANT2: ADP/ATP translocase 2
VDAC: voltage-dependent anion-selective channel protein 1
ATF6: activating transcription factor 6

## Author contributions

Study concept and design: S.Y. and S.S. Acquisition of data: S.Y., K.N., T.M., K.K., T.H., R.K., I.H., and S.S. Analysis and interpretation of data: S.Y., K.N., and S.S. Drafting of the manuscript: S.Y., T.H., and S.S. Obtained funding: S.Y., I.H., and S.S

## Acknowledgment

This work was funded by the Japan Society for the Promotion of Science grants-in-aid (KAKENHI) grant numbers 23K05996 (to S. S.) and 21K20701 (to S. Y.), the Research Support Project for Life Science and Drug Discovery (Basis for Supporting Innovative Drug Discovery and Life Science Research) from AMED under Grant Number JP21am0101120 (to I. H), Gout and Uric Acid Research Foundation 2020 (to S. S) and 2022 (to S. Y.), and Waseda University Grants for Special Research Projects 2021C-611 (to S. Y.), 2022C-611 (to S. Y.), 2022C-214 (to S. S.), and 2023C-207 (to S. S.). We would like to thank Ms. M. Mori and Ms. M Hidaka for their technical assistance. We also thank Enago (www.enago.jp) for English editing.

## SUPPLEMENTAL INFORMATION

Fig. S1. Congo red and PAS staining in the *Nwd1*^−/−^ liver

Congo red staining of wild-type and *Nwd1*^−/−^ livers. Nuclei were counterstained with hematoxylin. (B) PAS staining with (lower) or without (upper) α-amylase in wild-type and *Nwd1*^−/−^ livers. (C) Quantitative comparison of the PAS^+^ area in wild-type and *Nwd1*^−/−^ livers. Welch’s *t*-test, **, P < 0.01. Scale bars, 50 (A) or 100 μm (B).

Fig. S2. The number of peripheral blood leukocytes was increased in *Nwd1*^−/−^ mice

(A) The relative counts of lymphocytes, neutrophils, eosinophils, and basophils in whole peripheral blood from wild-type or *Nwd1*^−/−^ adult mice. (B) The ratios (%) of lymphocytes, monocytes, neutrophils, eosinophils, and basophils between wild-type and *Nwd1*^−/−^ mice.

Fig. S3. HE staining of other organs in *Nwd1*^−/−^ mice

(A–G) HE staining of lung, heart, intestine, kidney, skeletal muscle, spleen, and testis tissue from wild-type and *Nwd1*^−/−^ adult mice. Scale bars, 50 (A, C, and F) or 20 μm (B, D, E, and G).

Fig. S4. Autophagy lysosomal pathway dysfunction was not observed in *the Nwd1*^−/−^ liver

(A, B) *Nwd1*^−/−^ livers were immunostained with anti-LAMP1 (*red*) (A) or anti-LC3A/B (*green*) and anti-KDEL antibodies (*magenta*) (B). Scale bars, 50 μm.

Fig. S5. Validation of ER fractions in *the Nwd1*^−/−^ liver

(A) WB of the total lysate and ER fractions of the liver using anti-SERCA2, anti-SEIPIN, anti-KDEL, and anti-α-tubulin antibodies.

Tables S1–S3. Halo-Nwd1-binding proteins identified by proteomic analysis

Each protein entry is presented with the respective protein name, gene symbol, spectral count, and UniProt accession number.

