## supplemental Data for "Induction of NASH in the *Nwd1^−/−^* mouse liver via SERCA2-dependent endoplasmic reticulum stress"

Fig. S1

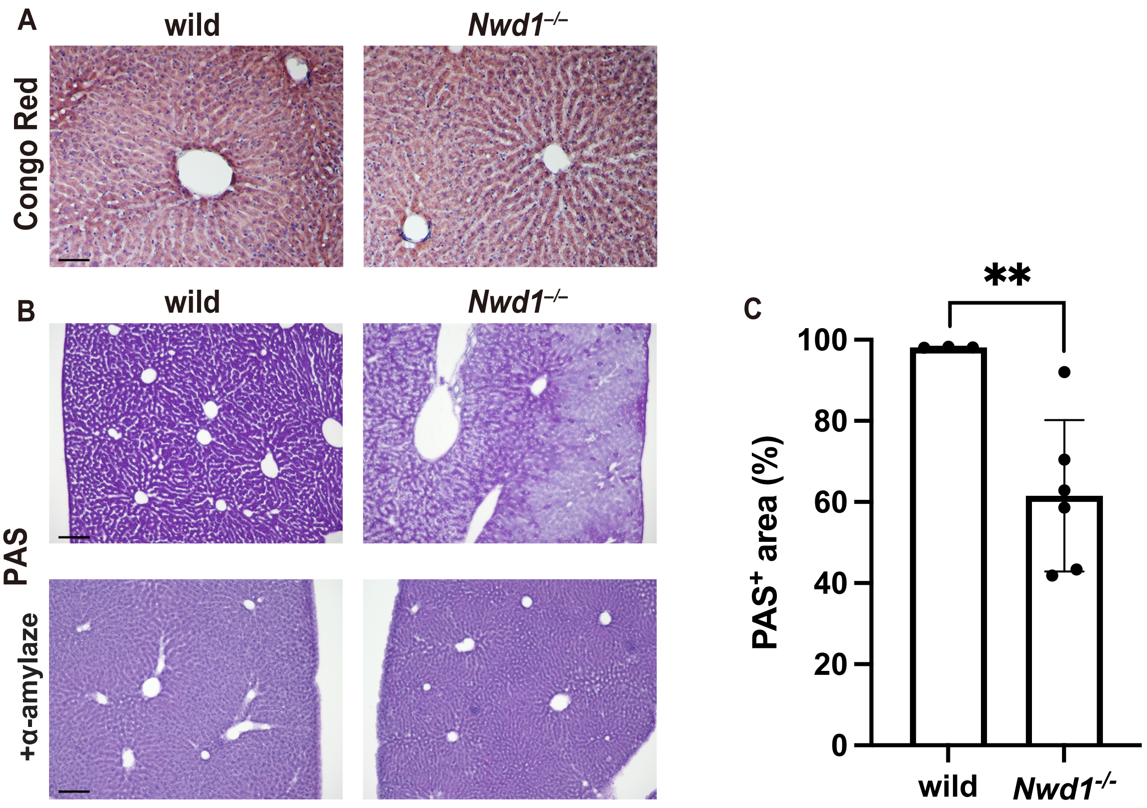

A
wild
Nwd1−/−
Congo Red
B
wild
Nwd1−/−
PAS
+α-amylaze
C
**
100
80
60
40
20
0
PAS+ area (%)
wild
Nwd1-/-

Fig. S2

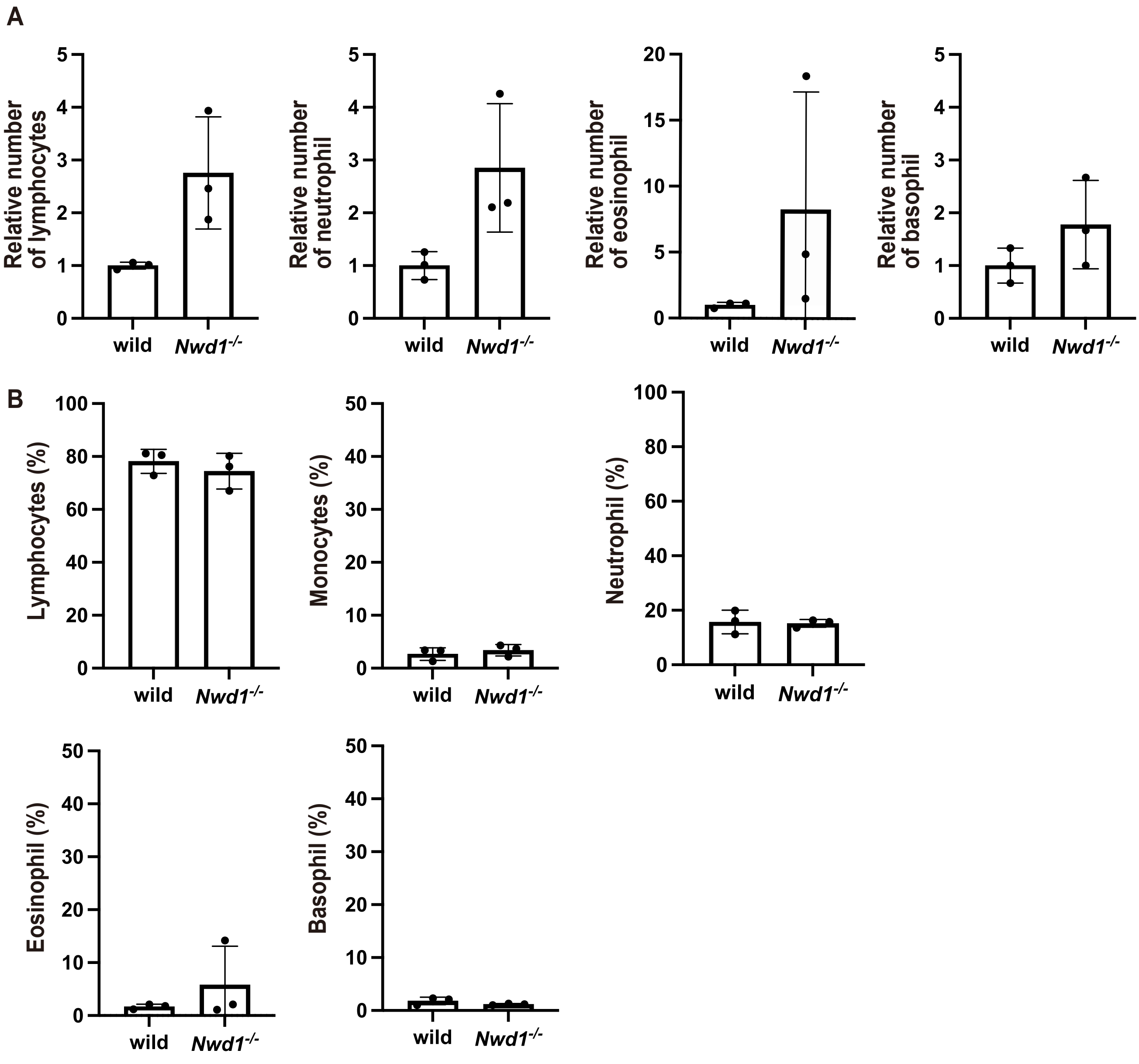

A
Relative number of lymphocytes
Relative number of neutrophil
Relative number of eosinophil
Relative number of basophil
wild
Nwd1-/-
B
Lymphocytes (%)
Monocytes (%)
Neutrophil (%)
Eosinophil (%)
Basophil (%)

Fig. S3

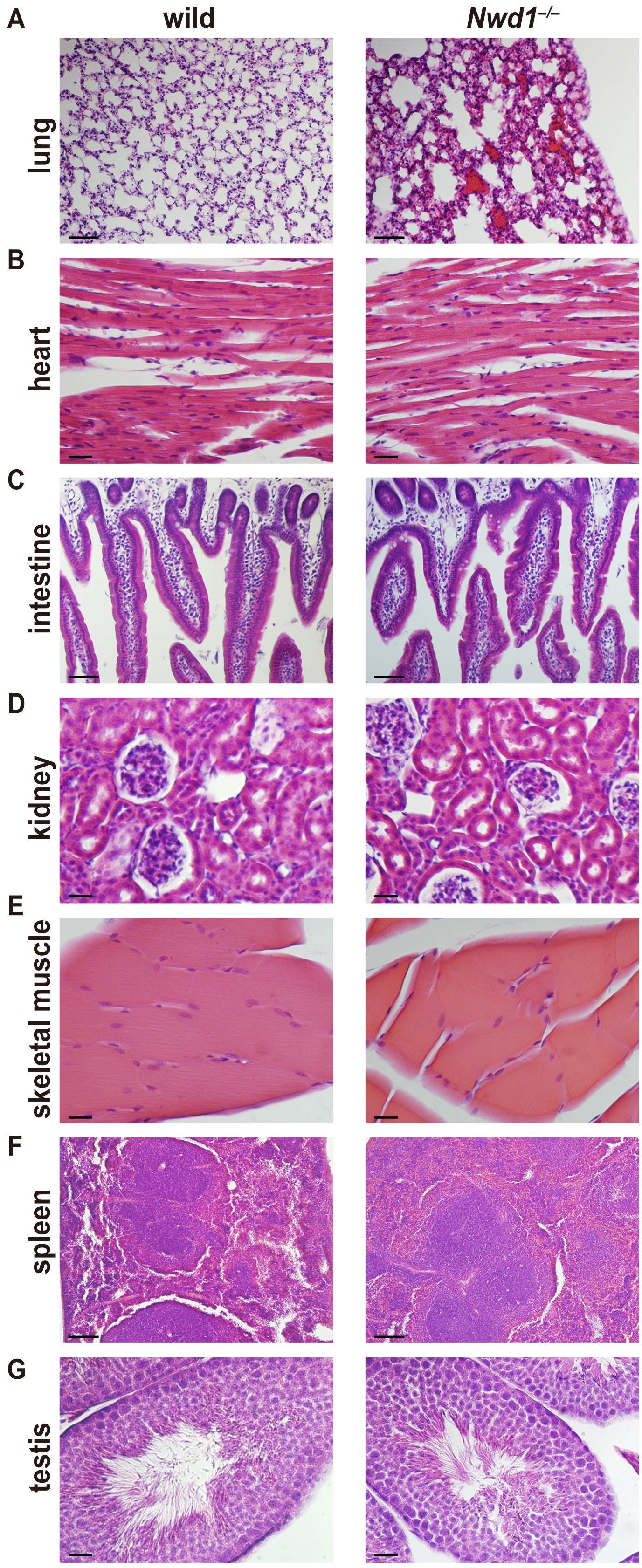
A
wild
$Nwd1^{-/-}$
lung
B
heart
C
intestine
D
kidney
E
skeletal muscle
F
spleen
G
testis

Fig. S4

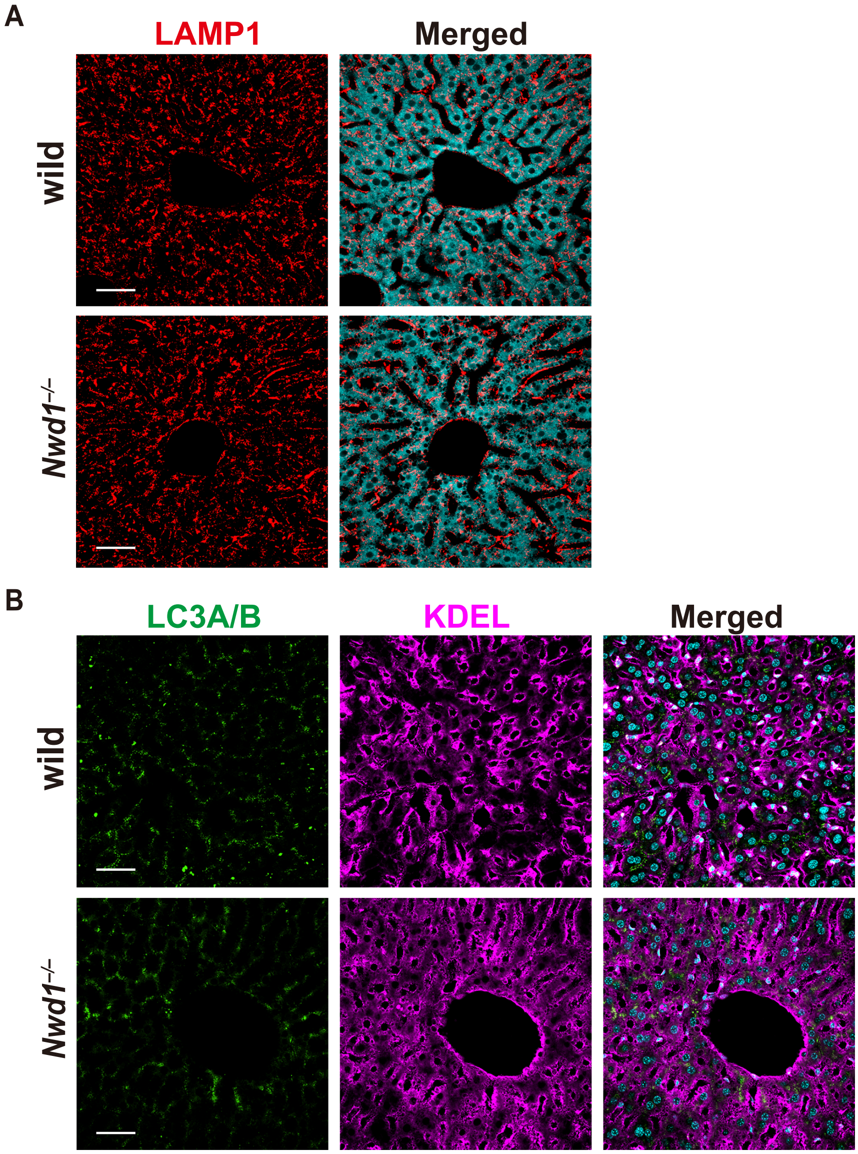

Fig. S5

A

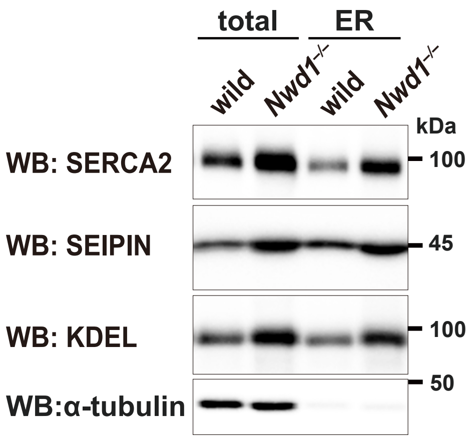

total
ER
wild
$Nwd1^{-/-}$
wild
$Nwd1^{-/-}$
kDa
WB: SERCA2
100
WB: SEIPIN
45
WB: KDEL
100
50
WB:α-tubulin

Table. S1

| Protein | Gene Symbol | Spectral Count | Accession |
|---|---|---|---|
| Sarcoplasmic/endoplasmic reticulum calcium ATPase 2 | ATP2A2<br>ATP2B<br>SERCA2 | 411 | P16615 |
| Tubulin beta chain | TUBB<br>TUBB5 | 400 | P07437 |
| Voltage-dependent anion-selective channel protein 1 | VDAC1<br>VDAC | 278 | P21796 |
| Calnexin | CANX | 261 | P27824 |
| Tubulin alpha-1B chain | TUBA1B | 258 | P68363 |
| Dolichyl-diphosphooligosaccharide--<br>protein glycosyltransferase 48 kDa subunit | DDOST<br>KIAA0115 | 229 | P39656 |
| Sideroflexin-1 | SFXN1 | 206 | Q9H9B4 |
| Dolichyl-diphosphooligosaccharide--<br>protein glycosyltransferase subunit 1 | RPN1 | 199 | P04843 |
| NADH-cytochrome b5 reductase 3 | CYB5R3<br>DIA1 | 197 | P00387 |
| Sodium/potassium-transporting ATPase subunit alpha-1 | ATP1A1 | 189 | P05023 |
| DNA-dependent protein kinase catalytic subunit | PRKDC<br>HYRC | 182 | P78527 |
| Ras-related protein Rab-7a | RAB7A<br>RAB7 | 181 | P51149 |
| Prohibitin-2 | PHB2<br>BAP | 173 | Q99623 |
| Translocation protein SEC63 homolog | SEC63<br>SEC63L | 167 | Q9UGP8 |
| Very-long-chain (3R)-3-hydroxyacyl-CoA dehydratase 3 | HACD3<br>BIND1 | 167 | Q9P035 |
| Heat shock 70 kDa protein 1A | HSPA1A<br>HSP72 | 164 | P0DMV8 |
| Membrane-associated progesterone receptor component 1 | PGRMC1<br>HPR6.6 | 160 | O00264 |
| Elongation factor 1-alpha 1 | EEF1A1<br>EEF1A | 135 | P68104 |
| Transmembrane emp24 domain-containing protein 10 | TMED10<br>TMP21 | 131 | P49755 |
| B-cell receptor-associated protein 31 | BCAP31<br>BAP31 | 126 | P51572 |
| NPC intracellular cholesterol transporter 1 | NPC1 | 123 | O15118 |
| Heat shock cognate 71 kDa protein | HSPA8<br>HSC70 | 123 | P11142 |
| Voltage-dependent anion-selective channel protein 3 | VDAC3 | 120 | Q9Y277 |
| Membrane-associated progesterone receptor component 2 | PGRMC2<br>DG6 | 120 | O15173 |
| CDGSH iron-sulfur domain-containing protein 2 | CISD2<br>CDGSH2 | 118 | Q8N5K1 |
| Voltage-dependent anion-selective channel protein 2 | VDAC2 | 114 | P45880 |
| CAAX prenyl protease 1 homolog | ZMPSTE24<br>FACE1 | 110 | O75844 |
| Vesicle-trafficking protein SEC22b | SEC22B<br>SEC22L1 | 110 | O75396 |
| Nodal modulator 1 | NOMO1<br>PM5 | 107 | Q15155 |
| Ras-related protein Rab-10 | RAB10 | 106 | P61026 |
| zfactb | LMAN1<br>ERGIC53 | 96 | P49257 |
| Phosphate carrier protein, mitochondrial | SLC25A3<br>PHC | 86 | Q00325 |
| Transmembrane emp24 domain-containing protein 5 | TMED5 | 86 | Q9Y3A6 |
| Basigin | BSG | 84 | P35613 |
| Cytochrome b-c1 complex subunit 2, mitochondrial | UQCRC2 | 80 | P22695 |

Table. S2

| Protein | Gene Symbol | Spectral Count | Accession |
|---|---|---|---|
| Transmembrane protein 33 | TMEM33<br>DB83 | 80 | P57088 |
| Cation-independent mannose-6-phosphate receptor | IGF2R<br>MPRI | 79 | P11717 |
| Transmembrane emp24 domain-containing protein 9 | TMED9<br>GP25L2 | 79 | Q9BVK6 |
| Cytochrome b5 type B | CYB5B<br>CYB5M | 79 | O43169 |
| CAD protein | CAD | 78 | P27708 |
| 7-dehydrocholesterol reductase | DHCR7<br>D7SR | 78 | Q9UBM7 |
| Cell division control protein 42 homolog | CDC42 | 77 | P60953 |
| NADPH--cytochrome P450 reductase | POR<br>CYPOR | 76 | P16435 |
| Mitochondrial import receptor subunit TOM70 | TOMM70<br>KIAA0719 | 76 | O94826 |
| Tubulin alpha-1C chain | TUBA1C<br>TUBA6 | 75 | Q9BQE3 |
| Endoplasmic reticulum-Golgi intermediate compartment-protein 1 | ERGIC1<br>ERGIC32 | 74 | Q969X5 |
| ADP/ATP translocase 2 | SLC25A5<br>AAC2 | 73 | P05141 |
| Dolichyl-diphosphooligosaccharide--protein glycosyltransferase subunit 2 | RPN2 | 71 | P04844 |
| Tyrosine-protein phosphatase non-receptor type 1 | PTPN1<br>PTP1B | 70 | P18031 |
| Alpha-soluble NSF attachment protein | NAPA<br>SNAPA | 69 | P54920 |
| Nicastrin | NCSTN<br>KIAA0253 | 67 | Q92542 |
| Transmembrane 9 superfamily member 4 | TM9SF4<br>KIAA0255 | 66 | Q92544 |
| Integral membrane protein GPR180 | GPR180<br>ITR | 66 | Q86V85 |
| Minor histocompatibility antigen H13 | HM13<br>H13 | 65 | Q8TCT9 |
| Torsin-1A-interacting protein 1 | TOR1AIP1<br>LAP1 | 65 | Q5JTV8 |
| Vesicular integral-membrane protein VIP36 | LMAN2<br>C5orf8 | 64 | Q12907 |
| Actin, cytoplasmic 1 | ACTB | 62 | P60709 |
| 4F2 cell-surface antigen heavy chain | SLC3A2<br>MDU1 | 61 | P08195 |
| Endoplasmic reticulum metallopeptidase 1 | ERMP1<br>FXNA | 61 | Q7Z2K6 |
| Dolichol-phosphate mannosyltransferase subunit 1 | DPM1 | 61 | O60762 |
| Transmembrane 9 superfamily member 3 | TM9SF3<br>SMBP | 60 | Q9HD45 |
| Transmembrane emp24 domain-containing protein 2 | TMED2<br>RNP24 | 60 | Q15363 |
| Transferrin receptor protein 1 | TFRC | 59 | P02786 |
| Autophagy-related protein 9A | ATG9A<br>APG9L1 | 59 | Q7Z3C6 |
| Syntaxin-16 | STX16 | 59 | O14662 |
| Translocon-associated protein subunit alpha | SSR1<br>TRAPA | 59 | P43307 |
| Cytochrome b-c1 complex subunit 1, mitochondrial | QCR1 | 58 | P31930 |
| Very-long-chain enoyl-CoA reductase | TECR<br>GPSN2 | 58 | Q9NZ01 |
| ER membrane protein complex subunit 1 | EMC1<br>KIAA0090 | 57 | Q8N766 |
| Syntaxin-binding protein 3 | STXBP3 | 56 | O00186 |

Table. S3

| Protein | Gene Symbol | Spectral Count | Accession |
|---|---|---|---|
| Heme oxygenase 2 | HMOX2<br>HO2 | 56 | P30519 |
| Mitochondrial carrier homolog 2 | MTCH2<br>MIMP | 56 | Q9Y6C9 |
| Oligosaccharyltransferase complex subunit OSTC | OSTC | 56 | Q9NRP0 |
| NACHT domain- and WD repeat-containing protein 1 | NWD1 | 54 | Q149M9 |
| Transmembrane protein 205 | TMEM205 | 54 | Q6UW68 |
| ATP synthase subunit alpha, mitochondrial | ATP5F1A<br>ATP5A | 53 | P25705 |
| Ras-related protein Rab-11B | RAB11B<br>YPT3 | 53 | Q15907 |
| PRA1 family protein 2 | PRAF2 | 51 | O60831 |
| Translocon-associated protein subunit delta | SSR4<br>TRAPD | 51 | P51571 |
| Inactive tyrosine-protein kinase 7 | PTK7<br>CCK4 | 49 | Q13308 |
| Polypeptide N-acetylgalactosaminyltransferase 2 | GALNT2 | 49 | Q10471 |
| Vesicle-associated membrane protein-associated protein B/C | VAPB | 49 | O95292 |
| Transmembrane emp24 domain-containing protein 1 | TMED1<br>IL1RL1L | 49 | Q13445 |
| Eukaryotic peptide chain release factor GTP-binding subunit ERF3A | GSPT1<br>ERF3A | 48 | P15170 |
| Calcium load-activated calcium channel | TMCO1<br>TMCC4 | 48 | Q9UM00 |
| Synaptophysin-like protein 1 | SYPL1<br>SYPL | 47 | Q16563 |
| Protein jagunal homolog 1 | JAGN1 | 47 | Q8N5M9 |
| GPI-anchor transamidase | PIGK<br>GPI8 | 46 | Q92643 |
| Signal peptidase complex catalytic subunit SEC11A | SEC11A<br>SEC11L1 | 46 | P67812 |
| Peroxisomal membrane protein 2 | PXMP2<br>PMP22 | 46 | Q9NR77 |
| Acylglycerol kinase, mitochondrial | AGK<br>MULK | 45 | Q53H12 |
| Ras-related protein Ral-A | RALA<br>RAL | 45 | P11233 |
| Ras-related protein Rap-1A | RAP1A<br>KREV1 | 45 | P62834 |
| Plasma membrane calcium-transporting ATPase 2 | ATP2B2<br>PMCA2 | 43 | Q01814 |
| Syntaxin-4 | STX4<br>STX4A | 42 | Q12846 |
| Vesicle-associated membrane protein 7 | VAMP7<br>SYBL1 | 42 | P51809 |
| Sodium bicarbonate cotransporter 3 | SLC4A7<br>BT | 41 | Q9Y6M7 |
| Peptidyl-tRNA hydrolase 2, mitochondrial | PTRH2<br>BIT1 | 41 | Q9Y3E5 |
| Transforming protein RhoA | RHOA<br>ARH12 | 40 | P61586 |
| Dolichyl pyrophosphate Man9GlcNAc2 alpha-1,3-glucosyltransferase | ALG6 | 39 | Q9Y672 |
| Ras-related protein Rab-2A | RAB2A<br>RAB2 | 38 | P61019 |
| Vitamin K epoxide reductase complex subunit 1-like protein 1 | VKORC1L1 | 38 | Q8N0U8 |
| Derlin-1 | DERL1<br>DER1 | 38 | Q9BUN8 |
| Monocarboxylate transporter 1 | SLC16A1 | 36 | P53985 |
| Transmembrane emp24 domain-containing protein 7 | TMED7 | 36 | Q9Y3B3 |
| Solute carrier family 35 member F6 | SLC35F6 | 36 | Q8N357 |
